# Lymphocytes and monocytes undergo swift suppression of IL-10R, IL-6R, and IL-2Rβγ signaling under high concentrations of different cytokines

**DOI:** 10.1101/2025.05.02.651967

**Authors:** Maxim Kuznetsov, Joao Rodrigues Lima Junior, Russell C. Rockne, Sergio Branciamore, Andrei S. Rodin, Peter P. Lee

## Abstract

The JAK-STAT signaling pathway is fundamental for immune system regulation. It involves phosphorylation of several types of STAT proteins in response to binding of cytokines to immune cell receptors. Traditionally, the immune signaling studies focus on measuring the levels of phosphorylated STATs (pSTATs) following individual cytokine application. We developed an experimental approach, based on multiparametric flow cytometry, to simultaneously measure the levels of five pSTATs after 15 minutes of cell treatment with high doses of individual cytokines and their paired combinations.

Analysis of our experimental data involving peripheral blood mononuclear cells from healthy donors reveals systematic suppression of IL-10R, IL-6R, and IL-2Rβγ signaling in T cells, B cells, NK cells, and monocytes. This suppression is mediated by at least all tested cytokines that do not induce relevant pSTATs by themselves. Remarkably, the cytokines with negligible own signaling do act as prominent selective signaling suppressors. In contrast, the signaling of IFNAR, IFNGR, IL-4R, and IL-2Rαβγ remains largely unaffected by co-application of other cytokines.

We propose that this pattern of signaling suppression represents an evolutionary developed mechanism enhancing the promptness, specialization, and efficiency of the immune response, while increased concentration of cytokines serves as a danger signal of inefficient response. We hypothesize that selective signaling suppression arises from the differential sensitivity of conformations of cytokine-receptor complexes to the increase of cell surface tension and stiffness, which is caused by effects following the binding of cytokines to membrane-associated molecules, including glycocalyx elements. While the rewiring of immune cell signaling should represent a powerful evolutionary tool for augmentation of adaptive response, it should also lead to the prolonged suppression of counteracting signaling pathways, culminating in cytokine release syndrome and contributing to autoimmune diseases.

## Introduction

One of the crucial components of immune cell signaling is the JAK-STAT pathway, which depends on interactions between Janus kinases (JAKs) with signal transducer and activator of transcription proteins (STATs) ^1^. JAK-STAT pathway promotes cell proliferation, differentiation, migration, and apoptosis, playing a key role in the immune system development and balance. In mammals, it serves as the primary signaling mechanism for numerous cytokines and growth factors ^2^. Their binding to different receptors, associated with common intracellular JAKs, alters receptor configurations bringing JAKs into close proximity. In this arrangement, JAKs phosphorylate tyrosine residues on each other and on the receptor, enabling STAT proteins to bind to the receptor and undergo phosphorylation. Phosphorylated STATs, or pSTATs, detach from the receptors, dimerize, and enter the nucleus to trigger transcription of target genes, leading to production of new receptors and cytokines, as well as to other downstream effects ^3^.

The JAK-STAT pathway is highly conserved across species from drosophila ^4^ to mammals, with structurally similar core components underscoring its evolutionary significance ^5,6^. Its signaling is modulated by a variety of regulatory proteins, including suppressors of cytokine signaling (SOCSs), protein inhibitors of activated STATs, and protein tyrosine phosphatases ^7^. Dysregulation of JAK-STAT signaling can result in various autoimmune diseases, leukemias, and weakened immune responses ^2,6,8^. Notably, during the recent COVID-19 pandemic, the involvement of JAK-STAT pathway was highlighted as a central regulator of hyperinflammation in critically ill patients ^9^.

The major significance of JAK-STAT pathway in immune cells promotes its experimental investigation, with relevant studies focusing both on the early stages of pathway activation by performing measurements of various pSTATs ^10–13^, and on the downstream functional outcomes of cytokine administration ^14–16^. Most of the studies on JAK-STAT signaling concentrate either on application of individual cytokines ^12,13,15,16^ or combined administration of cytokines without comparing the outcomes to individual cytokine effects ^10,11^. While several earlier studies explored in detail the phenomena resulting from specific pairs of co-administered cytokines ^17–20^, such research has become infrequent lately ^21^.

However, it is well known that in physiological conditions, cellular behavior is shaped by simultaneous and context-dependent action of multiple cytokines. Some cytokines can exert opposing pro-inflammatory and anti-inflammatory effects depending on cell type, timing, dosing, and interactions with other cytokines. Therefore, many cytokines are termed pleiotropic, and their effects are often described as ambiguous ^22–24^. Similarly, the immune system as a whole can be both protective and harmful to the host. These contradictory effects stem at least in part from the immune system’s need to recognize and respond to a wide variety of threats. To ensure an efficient response, evolution has shaped several interconnected yet inevitably conflicting strategies, each relying on JAK-STAT signaling. The three major types of immune responses are type 1 response, targeting intracellular microbes and abnormal host cells; type 2 response, protecting the body from parasites and allergens; and type 3 response, combating extracellular bacteria and fungi ^25^. While these strategies may incur severe side effects for certain individuals, the specialization and adaptability of immune system are of paramount importance for the overall survival of the species ^26^.

Motivated by the pleiotropism of cytokine roles, we developed an experimental approach using multiparametric flow cytometry to analyze the effects of individual cytokines and cytokine pairs on immune cells derived from the peripheral blood of healthy human donors (thus representing a healthy baseline immune signaling system). Our methodology enables simultaneous measurement of five STAT types, focusing on short-term immune cell responses with cytokine treatment limited to 15 minutes. This time frame minimizes the influence of newly transcribed cytokines and receptors on the experimental outcomes. Our experimental data reveals the systematic suppression of JAK-STAT signaling of certain cytokine-reeptor complexes in T cells, B cells, NK cells, and monocytes. In this paper, we dissect these suppression patterns, propose their evolutionary roles, and put forward a mechanistic hypothesis for the systematic signaling suppression.

## Results

### Immune cell pSTATome is cell type-specific

In the first experimental setting, we investigated the effects of eight individually applied cytokines on the phosphorylation levels of five STATs across immune cells in absence of activating factors: T cells (Figure 1A), monocytes (Figure 1B), B cells (Figure 1C), and natural killer (NK) cells (Figure 1D). The immune cells were obtained from nineteen donors, with an equal number of cells from each donor selected for every setting in the presented aggregate analysis. Cytokines were administered at 50 ng/mL dose each, except for IFN-α which dose was 1000 IU/mL. Cytokine effects were quantified using the Cumulative Shift Index (CSI), based on rank-biserial correlation for Mann–Whitney U test (see Materials and

**Figure 1.**
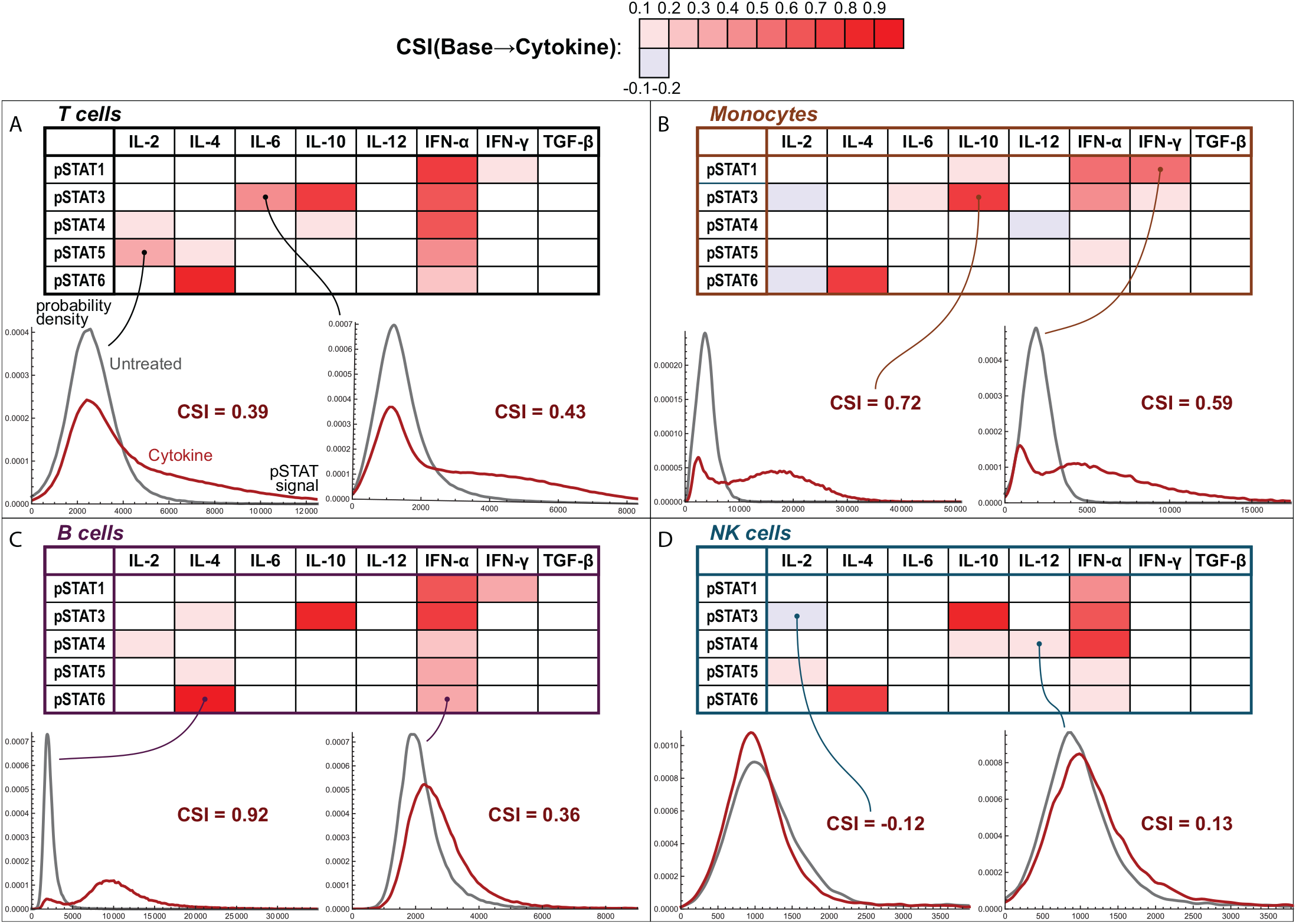
Effects of individual cytokines on STATs phosphorylation. Cytokines were applied for 15 minutes to the immune cells from nineteen healthy donors: **A)** T cells, **B)** monocytes, **C)** B cells, **D)** natural killer (NK) cells. Colored cells in the tables indicate significant stimulation or inhibition of specific pSTATs, with all corresponding p-values being less than 0.001 (Mann–Whitney U test). Effect strength is quantified by Cumulative Shift Index (CSI), based on rank-biserial correlation for Mann–Whitney U test. Distributions of pSTAT levels in untreated and treated cells are shown for eight specified settings. Here and further each cytokine was applied at 50 ng/mL dose, except for IFN-α which dose was 1000 IU/mL; cell counts are provided in the supplementary Excel (.xlsx) files.

Methods). The CSI value of 1 or -1 would correspond to non-overlapping distributions, while its values near 0 indicate minor shifts. To ensure robust analysis, we consider only the shifts with |CSI| ≥ 0.1 and p-values *<* 0.001 as significant.

The results consistently reproduce the canonical signaling axes. IL-10/STAT3 and IL-4/STAT6 signaling are prominent in all the considered cell types and their identified subtypes (Figure S1A), and for each donor (Figure S1B). In certain cell types, these dominant signaling axes are complemented by moderate phosphorylation of additional STATs. In particular, IL-10/STAT1 signaling is evident in classical and non-classical monocytes, as well as in memory B cells.

Less abundant signaling, varying across donors, and restricted to specific cell types, is induced by IFN-γ, IL-2, and IL-6. The primary IFN-γ effect is pSTAT1 signaling, not detected in resting NK cells; IL-2 induces STAT5 in T cells and NK cells; and IL-6 phosphorylates STAT3 in classical monocytes and T cells. In certain cell types, the signaling profiles of these cytokines are as well supplemented by moderate induction of other pSTATs.

IFN-α demonstrates the most diverse action. Our experimental panel did not include an antibody for pSTAT2, which has a narrow activity profile and functions primarily as a part of a heterodimer with pSTAT1^27^. Literature provides direct evidence that IFN-α phosphorylates STAT2 in T cells ^28^ and B cells ^29^. Thus, we can conclude that IFN-α induces phosphorylation of every STAT in adaptive immune cells. However, in monocytes it has no significant effect on STAT6, which is notably expressed in these cells, as evidenced by IL-4 signaling in them. Additionally, classical monocytes do not exhibit IFN-α/STAT4 signaling, which should stem from the negligible levels of STAT4 in resting monocytes ^30^.

IL-12 shows only weak activity within 15-minute treatment, likely due to the scarcity of functional IL-12 receptors in resting T cells, B cells ^31^, and monocytes ^30^. As a true negative control, TGF-β shows no significant signaling at the aggregate level, which is consistent with the lack of literature evidence supporting its involvement in JAK-STAT signaling of immune cells ^12^. We have previously shown that TGF-β also induces minimal signaling through its canonical pathway in this experimental setting ^32^. This aligns with other reports showing modest TGF-β signaling and low expression of its signaling components in resting lymphocytes ^33,34^ and monocytes ^35^.

We observe several additional effects at the aggregate level, where cells display reduced pSTAT concentrations after treatment. Inhibition of pSTAT3 levels by IL-2 in NK cells and monocytes may result from the selective formation of pSTAT3:pSTAT5 heterodimers ^36^, interfering with pSTAT3 binding to the assay antibodies. Downregulation of pSTAT4 by IL-12 in classical monocytes is less noteworthy due to the negligible expression of both IL-12Rs and STAT4 in their basal state ^30^.

Overall, these results align with and expand upon the recent studies exploring short-term effects of individual cytokine applications in human immune cells across more limited cell type repertoires ^12,13^.

### IL-10, IL-6 and IL-2 signaling is systematically suppressed by other cytokines

Concurrently with the above experiments, we performed paired administration of the same doses of IL-10 in conjunction with other cytokines to the immune cells from the same donors. The results reveal notable suppression of IL-10/STAT3 signaling by different cytokines across all the considered cell types: T cells (Figure 2A), monocytes (Figure 2B), B cells (Figure 2C), and NK cells (Figure 2D), as well as in all of their identified subtypes (Figure S2A). Additionally, we observe suppression of IL-10/STAT1 signaling in both types of monocytes and memory B cells, where this axis is prominent. To quantify these effects, we introduce the Level of Suppression metric (LoS, see Materials and Methods), which takes values from 0 if no significant suppression is observed to 1 if the pSTAT distribution of cells treated with two cytokines matches that of untreated cells.

**Figure 2.**
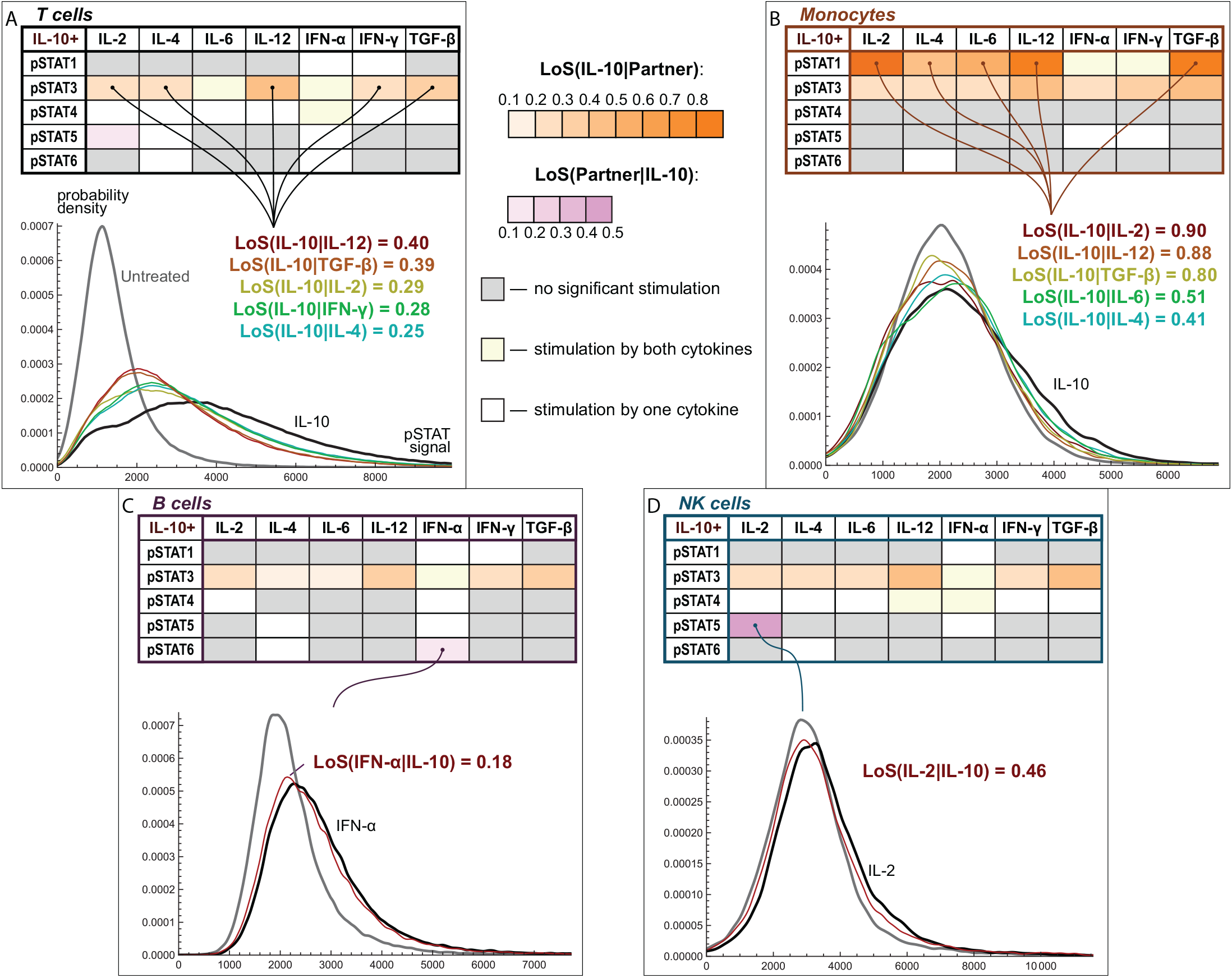
Effects of paired conjunction of IL-10 with other cytokines on STATs phosphorylation. Cytokines were applied for 15 minutes to the immune cells from nineteen healthy donors: **A)** T cells, **B)** monocytes, **C)** B cells, **D)** NK cells. Orange cells in the tables represent suppression of IL-10 signaling by a co-applied cytokine. Purple cells indicate suppression of a co-applied cytokine signaling by IL-10. The suppression of cytokine *a* by cytokine *b* is highlighted only for the cases where CSI(*Base → a*) ≥ 0.1 with *p <* 0.001 and − CSI(*a → ab*) ≥ 0.05 with *p <* 0.001. Level of Suppression (LoS) is evaluated as LoS(*a*|*b*) = − CSI(*a* → *ab*)*/*CSI(*Base → a*). Gray cells denote the settings where neither of two cytokines induces the relevant pSTAT significantly, both related p-values being ≥ 0.001. Yellow cells highlight the settings where both cytokines significantly induce the relevant pSTAT, interfering with potential detection of suppression. Distributions of pSTAT levels in untreated and treated cells are shown for four settings.

Detection of IL-10 signaling suppression by IFN-α and IL-6 is not always feasible, since a reduction in the levels of pSTAT3 can be compensated by its supplemental induction. Nevertheless, suppression of IL-10 signaling by IFN-α is evident in both types of monocytes and several T cell subsets, while suppressive action of IL-6 is detected in all cell types where IL-6/STAT3 signaling is negligible, and also in several cell types where it is prominent: classical monocytes, memory B cells, and several T cell subtypes.

Across all the studied settings, TGF-β and IL-12 emerge as the most potent suppressors of IL-10 activity, despite displaying negligible or weak JAK-STAT signaling by themselves. The level of suppression notably varies among donors, with some of them exhibiting weaker signaling suppression in most settings (Figure S2B).

Our results also show that IL-10 can interfere with the effects of other cytokines. IL-2/STAT5 signaling is significantly suppressed by IL-10 in almost all cells where this axis is prominent: T cells, the majority of their isolated subtypes, and NK cells. IFN-α/STAT6 signaling is slightly suppressed in naive and memory B cells, but the phosphorylation of other STATs induced by IFN-α remains unaffected by IL-10 across all studied cell types. In contrast, this aggregate analysis shows no evidence of IL-10-induced suppression of IL-4 and IFN-γ signaling.

These findings motivated us to conduct additional experiments focused on different cytokine pairs. The detailed aggregate and individual donor analyses for all cell types and their identified subtypes are provided in supplementary materials for combinations involving IL-2 (Figure S3), IL-6 (Figure S4), IL-4 (Figure S5), and IFN-α (Figure S6).

The induction of pSTAT5 by IL-2 in T cells and NK cells is suppressed by each co-administered cytokine, except IFN-α, which induces pSTAT5 by itself (Figure 3A). Co-application of IL-4 shows the weakest but statistically significant suppression of IL-2/STAT5 signaling in T cells, with CSI = − 0.04, which formally falls below our stringent detection threshold. In CD8+ TEMRA cells of two donors, IL-2 also induces pSTAT3, which is as well suppressed by all cytokines not signaling through it.

**Figure 3.**
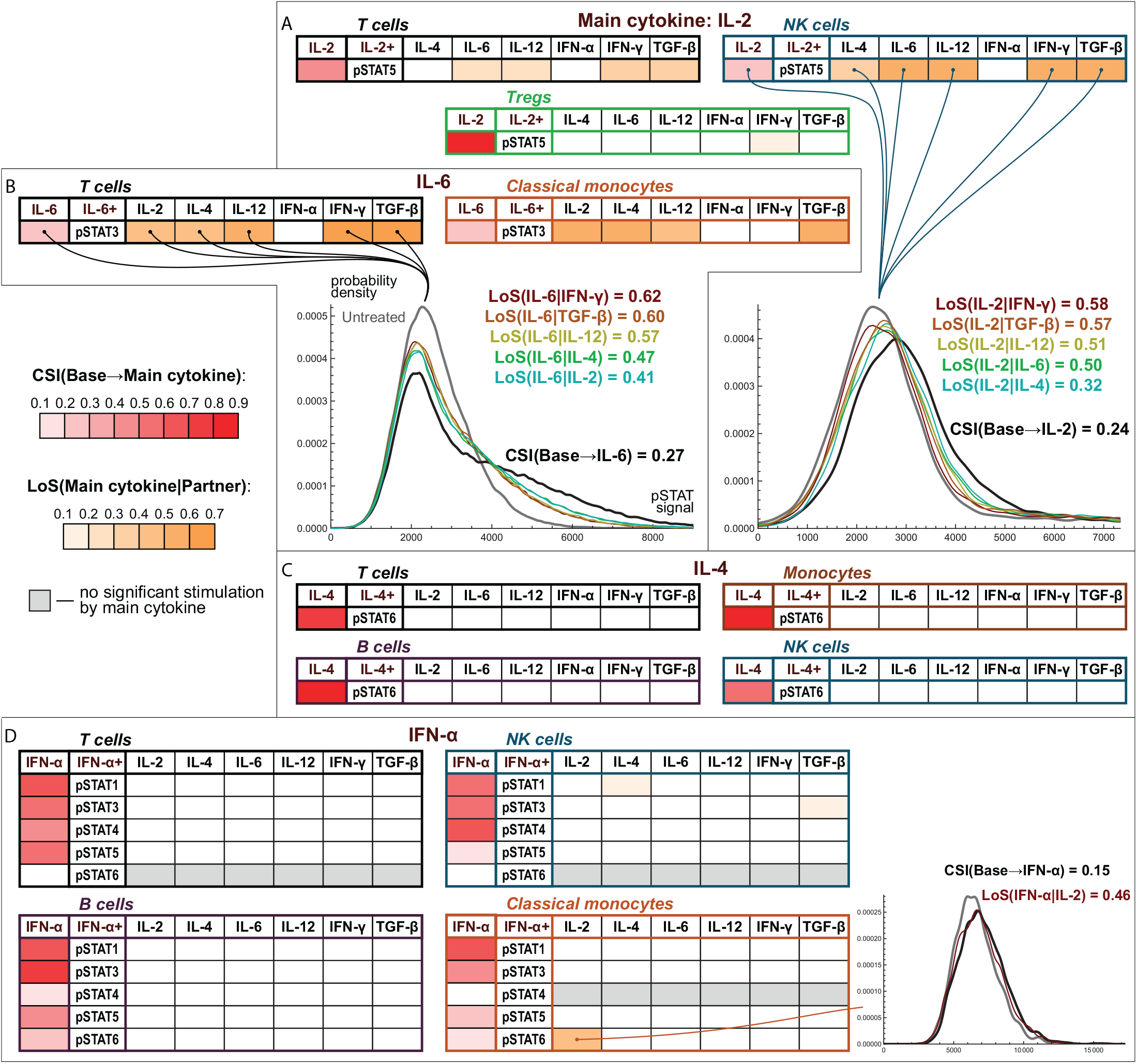
Effects of paired conjunction of cytokines on STATs phosphorylation. Cytokines were applied for 15 minutes to the immune cells from the groups of five healthy donors each: **A)** IL-2, **B)** IL-6, **C)** IL-4, **D)** IFN-α (main cytokines) in paired conjunction with other cytokines. Orange cells in the tables represent suppression of main cytokine signaling by a co-applied cytokine. The suppression of cytokine *a* by cytokine *b* is highlighted only for the cases where CSI(*Base* → *a*) ≥ 0.1 with *p <* 0.001 and − CSI(*a* → *ab*) ≥ 0.05 with *p <* 0.001. Gray cells denote the cases where the main cytokine does not induce the relevant pSTAT significantly (*p* ≥ 0.001). Distributions of pSTAT levels in untreated and treated cells are shown for three specified settings.

However, IL-2/STAT5 signaling is remarkably robust in each of the identified subtypes of regulatory T cells (Tregs). At the individual donor level, we detect only two related instances of suppression in Tregs. Intriguingly, resting T cells and NK cells express the intermediate-affinity form of IL-2 receptor (IL-2Rβγ), while Tregs are known to express its high-affinity form (IL-2Rαβγ) ^37^.

IL-6/STAT3 signaling in T cells and classical monocytes is suppressed by all co-applied cytokines, which do not induce pSTAT3 in corresponding cells (Figure 3B). This effect is consistently detected across all T cell subtypes that exhibit IL-6 signaling. In contrast, IL-4 induces pSTAT6 robustly in all considered cell types (Figure 3C). Only one donor exhibits consistent, but weak, suppression of IL-4/STAT6 signaling by each co-administered cytokine, except IFN-α, within a single cell type (NK cells). In this assay, we also performed co-treatment of cells with IL-10 and IL-4, which reproduced the previously observed outcome – IL-10 signaling was suppressed across all cell types, while IL-4 signaling was largely unaffected by IL-10 (Figure S5A).

The multifaceted signaling of IFN-α is largely resistant to the co-administered cytokines (Figure 3D). At the aggregate level, the only observed suppression events involve IFN-α/STAT6 axis in classical monocytes, naive B cells, and effector memory T cells; IFN-α/STAT1 axis in NK cells and CD8+ TEMRA cells; and IFN-α/STAT3 axis in NK cells.

### Suppressive power of cytokines rarely depends on their signaling level

Direct comparison of the signaling suppression levels shows that IL-10/STAT3 signaling undergoes stronger suppression by IFN-γ in monocytes than lymphocytes (Figure 4A). This correlates with the higher levels of IFN-γ/STAT1 signaling in monocytes and happens despite the fact that IFN-γ by itself significantly induces pSTAT3 in monocytes, complementing the action of IL-10. In contrast, for four other cytokines that do not typically induce pSTAT3, IL-10/STAT3 suppression levels are similar in monocytes and lymphocytes.

**Figure 4.**
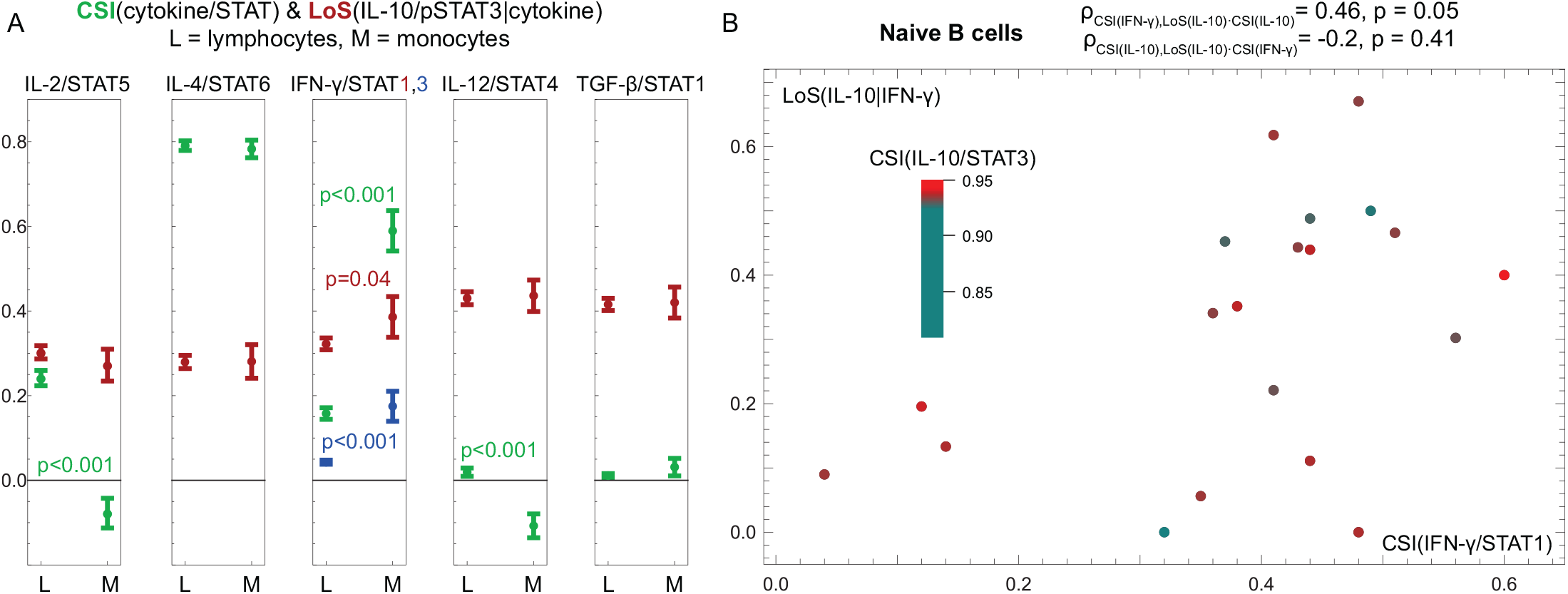
IFN-γ signaling correlates with IL-10/STAT3 signaling suppression. **A)** Comparison of signaling intensities (CSI) of IL-2, IL-4, IFN-γ, IL-12, and TGF-β alongside the levels of IL-10/STAT3 signaling suppression (LoS) in lymphocytes and monocytes. The average CSI and LoS values for twenty individual donors were aggregated from classical and non-classical monocytes (M) and eleven distinct types of lymphocytes (L): NK cells; memory and naive B cells; naive, central memory, TEMRA, and effector memory CD4+ T cells; and equivalent subtypes of CD8+ T cells. The p-values were estimated by the Mann–Whitney U test. **B)** Scatter plot of IFN-γ-induced suppression of IL-10/STAT3 signaling and the intensities of IFN-γ/STAT1 and IL-10/STAT3 signaling in naive B cells from twenty healthy donors. Coefficients of partial correlation between suppression levels and the intensity of each of the two axes while controlling for the intensity of another axis, *ρ*, and corresponding p-values were estimated by Spearman’s rank test. Among all the cell types and signaling axes of cytokines co-administered with IL-10 (IL-2/STAT5, IL-4/STAT6, IL-6/STAT3, IFN-γ/STAT1, and five IFN-α axes), this is the only setting showing a statistically significant partial correlation between the levels of co-applied cytokine signaling and IL-10/STAT3 suppression.

We also examined the relationships between the levels of cytokines’ signaling and IL-10/STAT3 suppression at the cell type-level using individual donor data. However, innate immune cells, and to a lesser extent CD4+ T cells, showed significant correlations between the signaling levels of different cytokines (with the notable exception of IL-2/STAT5 axis), which interfered with the task of isolating the separate influence of cytokine signaling on IL-10/STAT3 suppression (Table S1). In contrast, B cells and CD8+ T cells exhibited more heterogeneous signaling patterns, possibly reflecting their more specialized roles in the immune system.

Overall, IFN-γ emerged as the only cytokine with significant partial correlation between its signaling and induced IL-10/STAT3 suppression within a single cell type, naive B cells (Figure 4B). For all other cytokines, we found no evidence of correlation between their signaling intensities and suppressive capabilities.

## Discussion

### Common pro-inflammatory effects of IL-10 are unlikely due to its direct action

As our experimental data suggests, sufficiently elevated cytokine concentrations should lead to the observed pattern of rewiring of lymphocyte and monocyte signaling in humans. We hypothesize that this rewiring serves to mobilize immune resources and enhance the efficiency of immune response, while abnormally high cytokine levels serve as a “danger signal” of the currently inefficient response.

IL-10 is recognized as a key anti-inflammatory cytokine ^38^, one of its functions being reflected in its original name, “cytokine synthesis inhibitory factor” ^39^. Its functions include promotion of B cell apoptosis upon activation ^40^, inhibition of NK cell responses ^41^, and suppression of T cell activation which IL-10 can induce both directly, by mitigating T cell receptor signaling ^42^, and indirectly, by reducing the antigen-presenting capacity of monocytes ^43^ and preventing dendritic cell maturation ^44^ (Table S2^45–50^). Thus, the nature of effects induced by IL-10 strongly suggests that the suppression of its signaling should augment the immune response.

However, multiple *in vitro* ^51–57^ and *in vivo* ^58–63^ experimental studies, along with several clinical investigations ^64–68^, provide evidence of the immunostimulatory effects associated with the high-dose IL-10 administration (Table S3). In *in vitro* experiments, these effects emerge upon immune cell activation after hours or days of IL-10 pre-treatment and include enhanced T cell survival ^51^, increased production of cytotoxic mediators ^59^ and cytokines ^52^ by T cells, amplified B cell proliferation and antibody secretion ^53^, and heightened NK cell cytotoxicity ^54,55^. Several studies explicitly demonstrate the dose-dependent nature of these effects, with their intensity saturating within an IL-10 dose range of ∼ 30-1000 ng/mL ^53–55,59^. *In vivo* and clinical studies corroborate these findings, demonstrating enhanced activation of T cells and NK cells ^59,63,66,67^, and their intensified tumor infiltration ^58,59,62,66^ under high dose IL-10 administration. In accordance with these findings, during the recent COVID-19 pandemic, high IL-10 level was identified as a predictor of severe disease progression ^69^, associated with a high proportion of hyperactivated T cells in the peripheral blood ^70^.

Our findings challenge the prevailing notion that these phenomena represent direct pro-inflammatory effects of IL-10^71^. Recently, Islam et al. ^72^ proposed that the inability of IL-10 to exert its anti-inflammatory effects during a cytokine storm may arise from the resistance or hyporesponsiveness of activated immune cells to it. Our results support this hypothesis and further suggest that the immune cell hyporesponsiveness to IL-10 represents a direct consequence of elevated cytokine levels. Notably, experimental data of Prince et al. ^73^ show that cytokine levels in tissues, when combined together, can reach more than 75 ng/g in a mouse model of chronic infection which, when benchmarked against our dosing regime, highlights the feasibility of selective signaling suppression in immune cells within inflamed tissues.

We further hypothesize that the aforementioned immunostimulatory effects are not only facilitated by IL-10 signaling suppression but also represent a biomechanical response of immune cells to abnormally high cytokine levels. Currently, physical forces are increasingly recognized as important immunity modulators, with alterations in tissue mechanics being acknowledged as the danger signal alerting immune cells to potential injury or infection ^74^. In particular, a physiologically realistic increase of substrate stiffness enhances activation of NK cells ^75^ and macrophages ^76^, induces the accumulation of BCR molecules in the immunological synapses of B cells ^77^, and elevates T cell cytokine production ^78^ and proliferation rate, while lowering the threshold dose of antigen required to induce T cell effector responses ^79^. Thus, mechanosensing serves as an important part of immune cell activation process. Therefore, it seems plausible that sufficiently high cytokine levels may as well enhance immune cell responsiveness via biomechanical effects.

Recent studies highlight another immunostimulatory function of IL-10, which is the prevention of activation-induced T cell exhaustion ^80,81^. While this effect may not completely align with the classical paradigm of anti-inflammatory role of IL-10, it is consistent with its characterization as a suppressor of immune cell activity ^82^, which encompasses its primary functions under normal physiological concentrations (Table S2).

### Suppression of IL-6 signaling should facilitate adaptive immune response

IL-6 is a crucial driver of the acute immune response. In reaction to infections and tissue injuries, IL-6 is produced by macrophages ^83^ to stimulate secretion of acute phase proteins by the liver. These proteins include serum amyloid A, a key promoter of type 3 immune response against extracellular bacteria and fungi, eliminated by innate immune cells.

Another role of IL-6 is facilitating the transition of immune response to a chronic stage, where eradication of pathogens relies on adaptive immune cells. In particular, IL-6 induces CD4+ T cell secretion of IL-4^84^, which plays a pivotal role in orchestrating type 2 response against parasites and allergens. IL-4 promotes migration of eosinophils ^85^ that kill pathogenic targets and mediates recognition of targets by eosinophils via inducing B cell secretion of pathogen-specific IgE antibodies ^86^.

Upon another major type of threat – viral infection – various virus-sensing immune cells secrete IFN-α ^87^, which establishes the foundation for type 1 response against intracellular microbes. In particular, IFN-α activates dendritic cells (DCs) ^88^ and NK cells ^89^, enhances the antigen-presenting function of monocytes ^90^, lowers the activation threshold of B cells ^91^, and promotes proliferation and cytotoxicity of CD8+ T cells ^92^, the primary killers of infected cells. Furthermore, IFN-α induces T cell secretion of IFN-γ, which reinforces type 1 response by driving DC maturation ^93^, boosting CD8+ T cells cytotoxicity and motility ^94^, and promoting differentiation of monocytes into phagocytic macrophages ^95^.

The effects of IL-6, IL-4, IFN-α, and IFN-γ interact in complex ways, reflecting the evolutionarily shaped strategies of the immune system to tailor the distribution of its resources for recognizing and combating a specific type of pathogen while minimizing the damage to healthy cells (Table S2^96–113^). Some of the effects of different cytokines, such as the IL-6-induced transition to a chronic stage, have complementary nature. Intriguingly, the diverse JAK-STAT signaling of IFN-α, marked by strong induction of pSTAT1 (main factor activated by IFN-γ) and moderate phosphorylation of STAT6 (main factor activated by IL-4), suggests that IFN-α actually fosters an initial broad type of response to be further stratified by more specialized cytokines. Experimental findings of Eriksen et al. ^114^ support this idea, showing that IFN-α induces a time-dependent effect in human T cells, lowering the threshold for IL-4-mediated STAT6 activation during the first 6 hours of administration, and inhibiting IL-4 signaling at longer time frames.

Nevertheless, the effects of different cytokines often inevitably conflict with each other (Table S2). For example, IL-4 inhibits expansion and migration of neutrophils ^115^, interfering with their IL-6-mediated effector functions. IFN-γ enhances activation and cytotoxicity of macrophages ^116^, while IL-6 and IL-4 on contrary promote their regulatory polarization for the resolution of inflammation ^117,118^. Additionally, IL-6, IL-4, and IFN-γ drive different polarization of helper T cells ^119–121^ and B cells ^86^ to fine-tune their functions for specific types of response. Furthermore, IL-6 suppresses NK cell cytotoxicity ^122^ and DC maturation ^123^, and also diverts CD8+ T cells from their crucial cytotoxic role by promoting their differentiation into regulatory cells ^124^ and B helper CD8+ T cells ^125^.

Thus, despite the essential role of IL-6 in initiating the chronic stage of immune response, its effects can be largely detrimental to the established type 2 and type 1 responses, which combat pathogens not recognized by the innate immunity. This reasoning is supported by clinical data, as inhibition of IL-2 and IFN-γ production by IL-6 was identified as a potential factor mitigating immune system performance during viral infections ^126^. Experimental studies further demonstrate that IL-6 can decrease cytotoxicity of NK cells ^122^ and monocytes ^127^, restrain activation of CD8+ T cells ^128^, and inhibit type 1 response-associated differentiation of CD4+ T cells ^129^. Therefore, suppression of IL-6 signaling should represent an effective evolutionary mechanism for the optimization of adaptive immune response (Figure 5A).

**Figure 5.**
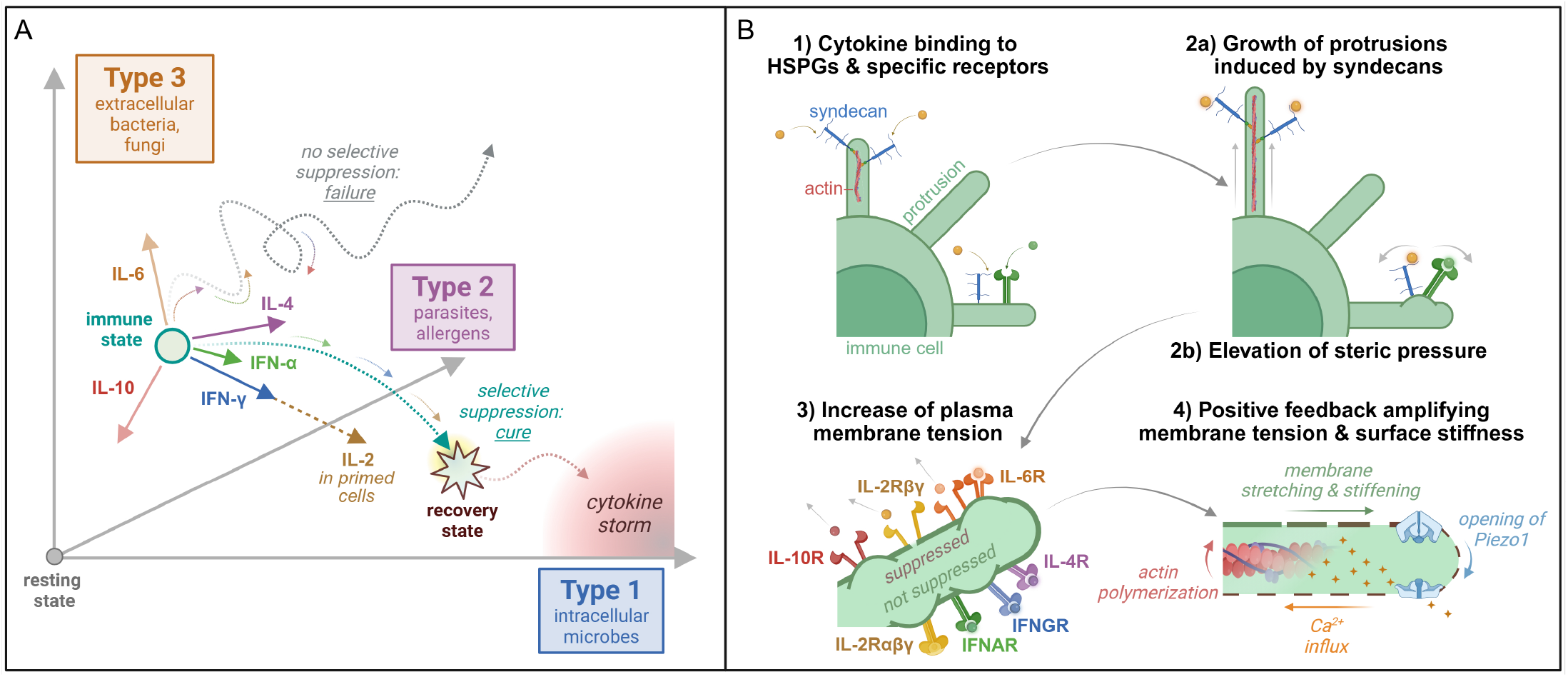
Hypothetical explanation of the observed selective suppression of cytokine signaling. **A)** Potential evolutionary reasons. Under abnormally high concentration of cytokines, serving as a danger signal of inefficient immune response, selective suppression of IL-10-induced inhibition of immune activity and IL-6-driven augmentation of type 3 response should enable mobilization of immune resources towards adaptive response addressing a critical risk. Differential suppressibility of IL-2 receptor forms should allow alleviating proliferative competition between immune cells, fostering proliferation of antigen-experienced rather than naive cells. **B)** Potential biomechanical reasons. Non-specific binding of cytokines to syndecans may induce actin polymerization, while cytokine binding to any membrane-associated molecules should increase steric pressure generated by their crowding. Both effects would result in deformations of cell plasma membrane and in increase of its tension. Structures of cytokine–receptor complexes may govern their differential sensitivity to membrane tension increase and thus yield their selective disruption. This effect should be intensified by the positive feedback mechanism amplifying membrane tension and stiffness.

Consistent with our aforementioned hypothesis on the existence of immunostimulatory effects driven by abnormally elevated levels of various cytokines, several experimental studies with high doses of IL-6 and IL-4 show immunostimulatory outcomes, mainly associated with the augmentation of type 1 response. In particular, in mice, injection of 10 µg of IL-4 induced production of IFN-γ by NK cells within several hours ^130^. In *in vitro* studies, a 6-day treatment of NK cells with 250 ng/mL of IL-6 significantly enhanced their expression of activation antigens ^131^, a 4-day treatment of cytotoxic T-cells with 5-40 ng/mL of IL-6 promoted their proliferation and inhibited viral replication in a dose-dependent manner ^132^, and 24-hour treatment of monocytes with 50-100 ng/mL of IL-6 suppressed the efficacy of their type 3-associated response against fungal pathogens ^133^. Similarly, type 1 response-associated upregulation of IFN-γ production was reported in experimental and clinical studies involving high-dose IL-10 administration ^52,54,59,64,66,67^. These data overall suggest that under abnormally high cytokine levels immune cells not only undergo selective suppression of their signaling but also prioritize enhancement of type 1 response, aiming at rapidly proliferating microbes and thus requiring a swift mobilization of resources to address the critical risk.

### Transient suppression of IL-2 should optimize generation of specialized subpopulations

IL-2 is a crucial amplifier of lymphocyte activity (Table S2^134–137^). It drives clonal expansion and differentiation of T cells ^138^; stimulates proliferation, activation, and cytotoxicity of NK cells ^139,140^; and promotes B cell proliferation and differentiation ^141^. Additionally, IL-2 regulates the homeostasis of Tregs which prevent excessive immune reactions ^142^.

Multiple experimental studies demonstrate activating effects of IL-2 on monocytes which, as our results show, exhibit negligible or very weak IL-2/STAT5 signaling. These studies involve high concentrations of IL-2 and report immunostimulatory effects ^143–146^, some of which vanish at physiological IL-2 levels ^147–150^. Therefore, we propose that these effects also reflect the immune cell reaction to abnormally high concentrations of cytokines, rather than represent consequences of direct IL-2 signaling. Thus, our findings highlight the need for revisiting the IL-2-induced effects in monocytes, as well as in macrophages ^147,151,152^.

A distinctive feature of IL-2 signaling is the existence of two forms of its receptor that utilize the same signaling machinery but differ in their binding affinities to IL-2^153^. Resting lymphocytes mainly express intermediate-affinity IL-2Rβγ, while high-affinity IL-2Rαβγ ^154^ is prominent on activated T cells ^37^ and NK cells ^155^, as well as on certain subsets of activated B cells ^156^. The explanations rationalizing the presence of two IL-2R forms generally focus on fine-tuning the sensitivity of different cells to IL-2^157^ and on competition for IL-2 among receptors sharing the common gamma chain subunit ^37^. However, these explanations do not address the reasons why differential IL-2 signaling relies on the modification of receptor rather than the regulation of its expression, which is a more straightforward and common evolutionary strategy – in particular, used for the enhancement of IFN-γ signaling in immune cells during type 1 response ^158^.

The study by Waters et al. ^159^ aimed to explore potential differences in IL-2 signaling through its two receptor forms. For that purpose, activated human T cells were incubated with or without IL-2Rα antibody daclizumab. This antibody was used at concentrations achievable in blood under clinical dosing, which are 15-30 µg/mL ^160^, resulting in ≳10^6^ molecules of daclizumab per each of the initial ∼10^4^ T cells. The kinetics of IL-2-induced pSTAT5 production were measured over half an hour, demonstrating a similar signal transduction in both groups. However, starting on the fourth day of incubation, daclizumab-treated T cells exhibited greater expansion driven by enhanced survival under maintained proliferation rate. A mathematical modeling study further suggested a more pronounced transmission of a secondary signal by IL-2Rβγ as the most plausible explanation for the observed effect.

To the best of our knowledge, direct evidence of differential induction of additional pathways by IL-2 through its two receptor forms is lacking. Given the aforementioned prominent evidence that abnormally high cytokine levels should enhance immune cell responsiveness, we hypothesize that the outcome of the experiment by Waters et al. might be caused by a similar biomechanically-driven reaction of immune cells to a high concentration of daclizumab. Notably, a similar effect of enhanced T cell survival despite a reduced proliferation rate was observed by Wang et al. ^51^ during a 3-day treatment of T cells with 1 µg/mL of IL-10.

The results of our study suggest that the existence of two IL-2R forms can be explained by the suppressibility of signaling through only one of them. Notably, IL-15R, the evolutionary precursor of IL-2R, appeared in jawed vertebrates (a group including sharks) and has a three-unit structure similar to that of the high-affinity IL-2Rαβγ. The divergence of IL-15R and IL-2R occurred in bony fish, with IL-15 still requiring the relevant α subunit for signaling, and IL-2 gaining the ability to signal through both three-unit and two-unit receptor forms ^161^.

This evolutionary route suggests that the existence of a suppressible IL-2R form confers an evolutionary advantage. This advantage may stem from the suppression of naive T cell proliferation (which can be promoted by monocytes in presence of IL-2^162^) during T cell recruitment to highly infected sites with elevated cytokine levels. The resulting alleviation of proliferative competition should further promote the expansion of antigen-experienced T cells expressing IL-2Rαβγ, thus enhancing the generation of effective responders under severe inflammation. A similar reasoning may also apply to NK cells, capable of developing antigen-specific memory ^163^.

### Structures of cytokine–receptor complexes may govern differential signaling suppressibility

Several earlier studies demonstrated that a short pre-treatment^19^ of pre-activated human ^17,20^ and murine ^18,19^ T cells with TGF-β, as well as its paired co-administration ^17,18,20^ with IL-2^17,18^ or IL-12^19,20^, suppresses the signaling of these interleukins within 10-20 minutes by inhibiting JAK phosphorylation. The mechanisms behind these effects were not identified, with proposed explanations including TGF-β-induced activation of some phosphatase ^19,20^ and allosteric effects ^19^. Additionally, recent experimental data by Cheemalavagu et al. ^21^ demonstrate decreased intensity of IL-10 signaling within the first 15 minutes of its co-administration with IL-6 in murine macrophages. Subsequent mathematical modeling reproduced this effect *in silico* under the assumption that IL-6-mediated transcription of SOCS1 inhibits phosphorylation of JAK1, associated with both IL-10R and IL-6R, in selective manner ^164^.

These hypotheses are not congruent with the signaling suppression pattern observed in our data, in particular with the effects mediated by cytokines with negligible signaling activity. Given the distinct stratification of cytokine-receptor complexes into suppressible and insuppressible groups, we hypothesize that the observed signaling suppression pattern arises from the differential sensitivities of membrane-embedded cytokine-receptor complexes to cell membrane tension. At a conceptual level, the specific structures of relevant receptors support our hypothesis.

IL-10 and IFN-γ receptor complexes are topologically similar, each forming a hexameric structure with a dimerized cytokine initiating signaling by engaging two separate dimerized receptor units ^165,166^. However, the IL-10 receptor complex has a nearly twice greater distance between these units, resulting in a significantly flatter profile compared to the IFN-γ receptor complex ^166^. This structural distinction suggests that IL-10R signaling may be more sensitive to increased membrane tension that tends to separate the receptor units and their associated JAK kinases.

Another hexameric receptor complex of IL-6 requires two ligand monomers for signaling ^167^. This feature may render its signaling more prone to disruption, as the dissociation of either monomer will also detach it from the signaling complex. The structure of IFNAR is marked by the significant flexibility of its low-affinity subunit, enabling it to optimize binding with the cytokine while being embedded in a moderately dynamic membrane. This feature was demonstrated in the molecular dynamics study by Li et al. ^168^, where the investigators proposed that the ability to probe multiple conformations should allow for the regulation of IFNAR signaling. Consistent with this notion, there exist multiple IFN-α subtypes, all of which, along with IFN-β and three other IFNs, bind to IFNAR ^169^ inducing quantitatively different broad-range STAT signaling and leading to different downstream effects ^170^ and functional outcomes ^171^. The intrinsic flexibility may also enable IFNAR to support and adjust its signaling in response to the changes in membrane tension, potentially explaining the selective suppression of certain axes, which emerges in our results.

IL-2Rβγ and both types of IL-4R are structurally similar, each consisting of two subunits. The binding affinity of IL-4 to its receptors is primarily governed by its interaction with IL-4Rα, which is present in both types of IL-4R ^172^. The dissociation constant for the binding of IL-4 and IL-4Rα is ∼ 1.5 · 10^−10^ M ^173^, while for the binding of IL-2 and IL-2Rβγ it is greater by nearly an order of magnitude ^174^. This difference may make IL-2Rβγ signaling less stable under elevated membrane tension compared to IL-4R signaling. The assembly of high-affinity IL-2R involves the IL-2Rα subunit, which captures IL-2 and presents it to the other subunits ^153^. This assembly decreases the relevant dissociation constant by two orders of magnitude, possibly rendering IL-2 signaling effectively insuppressible.

Our theoretical reasoning is also supported by the growing recognition of membrane geometry and tension as the key modulators of T cell and B cell antigen receptor signaling ^175^ and by the molecular dynamics modeling study of Simunovic and Voth ^176^, showing that elevated tension can inhibit interactions between certain membrane-bound proteins, impairing their assembly and reducing the lifetime of dimers. These results thus support the plausibility of a similar cell membrane involvement in the regulation of signaling of cytokines.

### Non-specific cytokine binding is the most plausible crucial driver of signaling suppression

Given the observed striking ability of cytokines with negligible signaling to suppress the signaling of other cytokines, we hypothesize that the cell membrane alterations, supposedly underlying this effect, are mainly induced by the non-specific binding of cytokines to membrane-associated molecules. Experimental studies show that the number of cytokine molecules non-specifically bound to immune cells initially grows linearly with increasing cytokine concentration ^177,178^. Weber-Nordt et al. ^179^ reported that non-specific binding of IL-10 in their experiments accounted for 2-3% of its total binding to murine B cells and Th1 cells, while saturation of IL-10R on 9*·*10^6^ cells of each type was achieved under administration of 7.5 ng of IL-10, of which 0.12 ng was bound. Given the IL-10 homodimer mass of ∼ 37 kDa ^180^, the linear extrapolation of these values to our setting (50 ng of IL-10 per 0.5-1 · 10^6^ cells) suggests that an average immune cell during our relevant experiments was non-specifically bound to ∼ 260-780 IL-10 molecules, and/or to a comparable number of other cytokine molecules, which is thus presumably sufficient to impact cell membrane dynamics.

Notably, all of the interleukins used in our experiments, along with IFN-γ and TGF-β, can bind either to heparan sulfate ^181–185^ or heparin ^186,187^, which infers that they interact with heparan sulfate proteoglycans (HSPGs) ^188^, the components of the glycocalyx layer covering the cell surfaces. HSPGs were shown to have profound effects on cell activity even at low expression levels ^189^, in particular, by promoting antigen presentation to T cells ^190^, regulating activation of T cells and NK cells ^191–193^, and controlling antigen uptake in B cells ^190^.

Notably, the cytoplasmic domains of syndecans, the members of HSPG family, are associated with the actin cortex and induce its rearrangement upon binding of syndecans to extracellular matrix ^194^. Syndecans can activate the signaling pathway that regulates actin polymerization within minutes even in response to soluble fibronectin fragments ^195^. It is therefore plausible that syndecan binding to cytokine molecules may as well trigger actin polymerization. Thus, syndecans may represent one of the key mediators for translating extracellular cytokine levels into mechanical effects.

Furthermore, syndecans were shown to induce the growth of microvilli ^196^, the finger-like protrusions facilitating lymphocyte interactions with antigens and other cells ^197–199^ and contributing to up to a half of the cell surface area ^200^. In line with our theoretical reasoning, Orbach et al. suggested that the deformations of microvilli may influence the organization of cytokine-receptor complexes and their activities ^201^. This suggestion can also be projected to the irregular ruffles of monocyte plasma membranes which, based on the estimates for macrophages, may constitute 20–40% of their cell surface area ^202^.

Intriguingly, the assembly of the IFN-γ receptor complex was shown to be largely facilitated by heparan sulfate oligosaccharides pulling this cytokine towards the cell surface ^203^. These active interactions may enhance cell membrane deformations induced by IFN-γ, explaining the observed correlation of the levels of IFN-γ/STAT1 signaling and IFN-γ-induced IL-10/STAT3 signaling suppression in some of our experimental settings.

In the absence of syndecan-induced actin polymerization, the binding of cytokines to HSPGs and other membrane-associated molecules can still contribute to the increase of cell membrane tension via elevation of the steric pressure arising from random collisions of crowded membrane-bound proteins, which was shown to be able to locally stretch and bend cell membranes regardless of the proteins’ structure ^204^. In particular, the increase of steric pressure may govern the observed suppression of IL-10/STAT3 signaling by IFN-α. IFN-α2a (the subtype used in our study) has a minimal binding affinity to heparin, suggesting that it does not interact significantly with HSPGs ^171^, however, the consistent JAK-STAT signaling of IFN-α across all considered cell types indicates that its receptors are ubiquitously expressed in them. Therefore, we hypothesize that for IFN-α the specific binding-related events accompanied by the significant flexibility of IFNAR ^168^ can drive bending and stretching of cell membranes and thus induce suppression of IL-10/STAT3 signaling, strong enough to compensate for the additional pSTAT3 induction by IFN-α.

The prominent sensitivity of immune cells to external cytokine levels can be explained by a positive feedback regulation

for the amplification of membrane tension, which was demonstrated by Atcha et al. ^76^. Mechanical stretching of cell plasma membranes triggers the opening of Piezo1 channels, which are present in lymphocytes and monocytes ^205^. In the open state, these channels allow intracellular influx of various cations, including Ca^2+^ ^206^. Among its effects, Ca^2+^ induces actin polymerization ^207^, leading to the stiffening of cellular cortex ^208^ as well as plasma membrane, which together form a composite cellular surface ^209^, and thus further promoting Piezo1-mediated Ca^2+^ influx. The actin cortex has a strain-stress relationship displaying notable non-linear growth starting from only ∼ 2-3% extensional strain. This indicates that its stiffness, i.e., resistance to deformation, grows with the increase of applied stress, i.e., force per cross-sectional area ^210^. Therefore, while a moderate increase of cell surface tension may disrupt the signaling of the assembled cytokine-receptor complexes, significantly high tension should also additionally enhance cell surface stiffness, thus inhibiting the assembly of certain complexes, as these processes rely on local membrane deformation ^211^.

The proposed set of hypotheses (Figure 5B) suggests that the extent of signaling suppression entailed by cytokines should depend on their size and shape, with larger and more structurally complex molecules expected to induce higher steric pressure and/or stronger syndecan-mediated actin polymerization. Intriguingly, our experimental data perfectly aligns with this reasoning, showing that large dimers with complex elongated shapes, IL-12 (with the mass of 70 kDa) and TGF-β (25 kDa), generally act as the most potent signaling suppressors despite the weak own signaling. In contrast, smaller and more compact monomers, IL-2 (15.5 kDa) and IL-4 ( ∼15 kDa) ^180^ demonstrate notably weaker suppressive capabilities.

### Potential implications for the immune system control

Overall, our theoretical reasoning highlights cell membrane as a potential major participant in the cytokine signaling processes, which enables rewiring of JAK-STAT signaling in lymphocytes and monocytes within a time frame of several minutes. Presumably, the differential sensitivity of cytokine receptors to membrane tension was shaped through gradual evolution, involving small changes in sensitivity and favoring the ones that optimize the overall immune response. Like any evolutionary shaped mechanism, it is plausible that the selective signaling suppression machinery is prone to errors in certain individuals ^32^, compromising the activation or deactivation of their immune systems. One of the drivers of such malfunction, and therefore a therapeutical target, may be cholesterol, with its content in a cell membrane significantly affecting its stiffness ^212^.

It is possible that other types of immune cells also leverage the mechanisms of selective signaling suppression, which may also extend to other cytokine-receptor complexes. This notion aligns with the experimental results of Houk et al. ^213^, who demonstrated that membrane tension acts as a long-range signaling inhibitor in migrating neutrophils, and with the findings of Pazdrak et al. ^214^, who showed that 15-minute pre-treatment of eosinophils with TGF-β suppresses their IL-5 signaling. It is also possible that selective signaling suppression can extend to non-immune cells, broadening the potential applications of the modulation of cell membrane composition as a therapeutic strategy ^215^. This notion is supported by the experimental results of Weber-Nordt et al. ^179^, who showed that 24-hour lipopolysaccharide treatment of fibroblasts reduced the affinity of their IL-10 receptors by more than an order of magnitude.

We hypothesize that, despite being a powerful evolutionary adaptation, selective signaling suppression machinery has a significant drawback, associated with the possibility of a prolonged suppression of IL-10 and IL-6 signaling, creating optimal conditions for triggering the cytokine release syndrome. The strategies aimed at restoring the immunosuppressive signaling of IL-10, such as generating its high-affinity mutants ^166,216^ and altering its shape and conformational transitions, can offer a promising approach to overcoming the hyporesponsiveness of immune cells to IL-10, and thereby alleviating cytokine storms. Such approaches would also address the lack of IL-10 efficacy in clinical settings for treating autoimmune and immune-mediated inflammatory diseases ^45,217^.

## Materials and Methods

### Peripheral blood samples

were collected from donors at the City of Hope Blood Donor Center, following the protocols approved by the Institutional Review Board (IRB 21368 and 19186). All donors were women with a mean age of 52.7 years. Blood was drawn into EDTA-treated tubes, and peripheral blood mononuclear cells (PBMCs) were isolated using Ficoll-Paque density gradient centrifugation (Cytiva, Marlborough, MA, USA), in accordance with the manufacturer’s instructions. The isolated PBMCs were then cryopreserved in a solution containing 10% dimethyl sulfoxide (DMSO) and fetal bovine serum (FBS).

### Cell culture

Cryopreserved PBMCs were thawed and incubated overnight during 16 hours in RPMI 1640 medium supplemented with 10% fetal bovine serum and 1% penicillin-streptomycin-glutamine (PSG) under controlled conditions: 37^*°*^C, 5% CO_2_. Cell counts and viability were assessed using a hemocytometer and the trypan blue exclusion method (Sigma-Aldrich). Following this, the cells were cultured in a 96-deep well plate at densities ranging from 0.5 to 1 × 10^6^ cells/mL in 200 µL of fresh RPMI 1640 medium (Thermo Fisher Scientific Inc., MA, USA).

### Cell signaling

After a resting period, PBMCs were cultured either untreated or stimulated at 37^*°*^C for 15 minutes with 50 ng/mL of IL-2, IL-4, IL-6, IL-10, IL-12, IFN-γ, or TGF-β, or 1000 IU/mL of IFN-α2a (PeproTech, Rocky Hill, NJ, USA), added to cultures using multi-channel pipette. The cells were then fixed with 25 µL 1.5% paraformaldehyde (PFA) for 10 minutes at room temperature. Following fixation, the cells were washed with 1x PBS and permeabilized using 500 µL of ice-cold 100% methanol. The methanol-fixed cells were subsequently stored at -80^*°*^C. Prior to antibody staining, the cells were washed three times with staining buffer (PBS supplemented with 1% FBS).

### Phospho-flow cytometry

was conducted using the following antibodies: Percp-Cy5.5 (clone 4a) for pSTAT1, AF488 (clone 4/P-Stat3) for pSTAT3, AF647 (clone 38/P-Stat4) for pSTAT4, STAT5-PE-Cy7 (clone 47), V450 (clone 18/P-Stat6) for pSTAT6, BV570 (clone UCHT1) for CD3, BUV563 (clone SK3) for CD4, BUV805 (clone SK1) for CD8, APC-Cy7 (clone HCD14) for CD14, BUV737 (clone 3G8) for CD16, AF700 (clone H1) for CD20, BV786 (clone L128) for CD27, BUV395 (clone HI100) for CD45RA, PE-CF594 (clone 259D/C7) for Foxp3. Antibody dilutions were prepared following the manufacturer’s guidelines and adjusted for optimal staining in preliminary experiments. Incubation was performed for 45 minutes at room temperature. All antibodies were obtained from BioLegend (San Diego, CA, USA) and BD Biosciences (Franklin Lakes, NJ, USA).

### Data acquisition and gating strategy

The stained cells were analyzed using a Cytek Aurora flow cytometer equipped with lasers operating at 355 nm, 405 nm, 488 nm, 561 nm, and 640 nm. Compensation settings were determined using single-stain controls and a negative control. Data acquisition occurred at a rate of 1,000 events per second, with each sample collecting between 50,000 and 100,000 events. Gating strategies for identifying cell populations are provided in supplementary materials (Figure S7).

The following cell types and subtypes were classified: T cells (CD3+), helper T cells (CD4+), cytotoxic T cells (CD8+), naive CD4+ T cells (CD4+CD45RA+CD27+), naive CD8+ T cells (CD8+CD45RA+CD27+), central memory CD4+ T cells (CD4+CD45RA-CD27+), central memory CD8+ T cells (CD8+CD45RA-CD27+), effector memory CD4+ T cells (CD4+CD45RA-CD27-), effector memory CD8+ T cells (CD8+CD45RA-CD27-), CD4+ TEMRA cells (CD4+CD45RA+CD27-), CD8+ TEMRA cells (CD8+CD45RA+CD27-), resting Tregs (CD4+CD45RA+FOXP3^*low*^), activated Tregs (CD4+CD45RA-FOXP3^*high*^), non-suppressive Tregs (CD4+CD45RA-FOXP3^*low*^), B cells (CD20+), memory B cells (CD20+CD27-), naive B cells (CD20+CD27+), NK cells (CD3-CD16+CD14-), monocytes (CD14+), classical monocytes (CD14+CD16-), and non-classical monocytes (CD14+CD16+).

### Mathematical methods

For evaluation of Cumulative Shift Index for a pair of cell groups *a* and *b*, CSI(*a* → *b*), we relied on rank-biserial correlation, which is a method to report the effect size for the Mann–Whitney U test. For its calculation, all the cells from both groups are sorted and ranked, with rank 1 assigned to the cell with the lowest level of pSTAT of interest, and the rank equal to the total number of cells in two groups *n*_*a*_ + *n*_*b*_ assigned to the cell with the highest level of that pSTAT. Then, Mann-Whitney U statistic for the group *b* is calculated as:

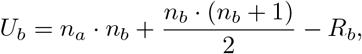

where *R*_*b*_ is the sum of ranks of elements in group *b*, and the first two terms represent its maximal possible value. Cumulative Shift Index is obtained by the normalization of *U*_*b*_:

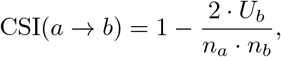

so that it can take values from -1 to 1. As follows from this definition, CSI(*a* → *b*) = − CSI(*b* → *a*).

For estimation of the Level of Suppression of the action of cytokine *a* under simultaneous application of cytokine *b*, LoS(*a*|*b*), we use the formula

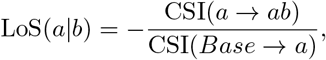

where *Base, a*, and *ab* represent the groups of untreated cells, cells treated only with cytokine *a*, and cells treated with both cytokines, respectively. Under this definition, a suppression that brings the distribution of pSTAT of interest within cells treated by two cytokines close to its basal distribution yields LoS(*a* |*b*) close to 1. For demonstration of the results, unless otherwise indicated, for individual cytokine administration we retain only the cases where |CSI(*Base* → *a*) | ≥ 0.1 with corresponding p-value estimated by the Mann–Whitney U test being *p <* 0.001, and for paired cytokine administration we retain only the cases where CSI(*Base* → *a*) ≥ 0.1 and − CSI(*a* → *ab*) ≥ 0.05 with both corresponding p-values being *p <* 0.001.

To ensure a balanced representation of each donor’s cells for the aggregate analysis, an equal number of cells per donor were selected randomly for each experimental setting. This selection was based on the minimum number of relevant cells available from any single donor for each experimental condition, involving the same cell type and the same cytokine application. Donor F67’s cells were excluded from the aggregate analyses, as they exhibited significantly lower basal pSTAT levels compared to other donors’ cells, potentially due to an underlying disease. For correlation analysis (Figure 4) and individual donor analyses, provided in the supplementary figures, all available cells from each donor were included.

### Code implementation

The computational codes were implemented in Wolfram Mathematica, version 14.1, and are provided in the supplementary files (https://doi.org/10.5281/zenodo.15330718), along with the raw experimental data, processed data arrays, and distribution plots for each of the experimental settings and correlation analysis. The provided data and codes ensure the reproducibility of the results, and also allow performing aggregate analyses with a random selection of an equal number of cells from each donor across all experimental settings.

## Disclosure of Potential Conflicts of Interest

The authors declare no potential conflicts of interest.

## Authors’ Contributions

P.P.L., A.S.R., S.B., and R.R. conceived the project. P.P.L., A.S.R., and R.R. obtained the funding. P.P.L., A.S.R., S.B., R.R., and M.K. designed the experiments. A.S.R. and P.P.L. supervised the project. J.R.L.J. performed the experiments. M.K. analyzed the data, revealed the systematic signaling suppression, and performed all the computational and theoretical work. A.S.R. and S.B. supervised the computational work. M.K. drafted the first version of the manuscript with input from A.S.R. and P.P.L., and M.K. wrote the final version with feedback from all co-authors.

## Acknowledgments

Research reported in this publication included work performed in the Analytical Cytometry and Biostatistics and Mathematical Oncology Shared Resources and was supported by the National Cancer Institute of the National Institutes of Health under grant numbers P30CA033572, U01CA232216 and by Dr. Susumu Ohno Chair in Theoretical Biology (held by A.S.R.). The content is solely the responsibility of the authors and does not necessarily represent the official views of the National Institutes of Health. The authors express their gratitude to the blood donors whose participation made this work possible and sincerely thank them for their contributions to the study.

## Notes

https://zenodo.org/records/15330718

### Competing Interest Statement

The authors have declared no competing interest.

## References

[1] George R Stark and James E Darnell. The JAK-STAT pathway at twenty. Immunity, 36(4):503–514, 2012.

[2] Jason S Rawlings, Kristin M Rosler, and Douglas A Harrison. The JAK/STAT signaling pathway. Journal of cell science, 117(8):1281–1283, 2004.

[3] Xiaoyi Hu, Jing Li, Maorong Fu, Xia Zhao, and Wei Wang. The JAK/STAT signaling pathway: from bench to clinic. Signal transduction and targeted therapy, 6(1):402, 2021.

[4] Natalia I Arbouzova and Martin P Zeidler. JAK/STAT signalling in Drosophila: insights into conserved regulatory and cellular functions. Development, 133 (14):2605–2616, 2006.

[5] Douglas A Harrison. The JAK/STAT pathway. Cold Spring Harbor perspectives in biology, 4(3):a011205, 2012.

[6] You Lv, Jianxun Qi, Jeffrey J Babon, Longxing Cao, Guohuang Fan, Jiajia Lang, Jin Zhang, Pengbing Mi, Bostjan Kobe, and Faming Wang. The JAK-STAT pathway: from structural biology to cytokine engineering. Signal Transduction and Targeted Therapy, 9(1):221, 2024.

[7] Chen Xue, Qinfan Yao, Xinyu Gu, Qingmiao Shi, Xin Yuan, Qingfei Chu, Zhengyi Bao, Juan Lu, and Lanjuan Li. Evolving cognition of the JAK-STAT signaling pathway: autoimmune disorders and cancer. Signal transduction and targeted therapy, 8(1):204, 2023.

[8] Rahul Pandey, Marina Bakay, and Hakon Hakonarson. SOCS-JAK-STAT inhibitors and SOCS mimetics as treatment options for autoimmune uveitis,psoriasis,lupus,and autoimmune encephalitis. Frontiers in immunology, 14:1271102, 2023.

[9] Ali Razaghi, Attila Szakos, Marwa Alouda, Béla Bozóky, Mikael Björnstedt, and Laszlo Szekely. Proteomic analysis of pleural effusions from COVID-19 deceased patients: Enhanced inflammatory markers. Diagnostics, 12(11):2789, 2022.

[10] Casper Marsman, Tineke Jorritsma, Anja Ten Brinke, and S Marieke van Ham. Flow cytometric methods for the detection of intracellular signaling proteins and transcription factors reveal heterogeneity in differentiating human B cell subsets. Cells, 9(12):2633, 2020.

[11] Wanxia Li Tsai, Laura Vian, Valentina Giudice, Jacqueline Kieltyka, Christine Liu, Victoria Fonseca, Nathalia Gazaniga, Shouguo Gao, Sachiko Kajigaya, Neal S Young, et al. High throughput pSTAT signaling profiling by fluorescent cell barcoding and computational analysis. Journal of immunological methods, 477:112667, 2020.

[12] Emily Monk, Melinda Vassallo, Paulo Burke, Jeffrey S Weber, Pratip Chattopadhyay, and David M Woods. Simultaneous assessment of eight phosphorylated STAT residues in T-cells by flow cytometry. bioRxiv, pages 2021–11, 2021.

[13] Leonard Daniël Samson, Peter Engelfriet, WM Monique Verschuren, H Susan J Picavet, José A Ferreira, Maryl`ene de Zeeuw-Brouwer, Anne-Marie Buisman, and A Mieke H Boots. Impaired JAK-STAT pathway signaling in leukocytes of the frail elderly. Immunity&Ageing, 19(1):5, 2022.

[14] Jia Xue, Susanne V Schmidt, Jil Sander, Astrid Draffehn, Wolfgang Krebs, Inga Quester, Dominic De Nardo, Trupti D Gohel, Martina Emde, Lisa Schmidleithner, et al. Transcriptome-based network analysis reveals a spectrum model of human macrophage activation. Immunity, 40(2):274–288, 2014.

[15] Sara Mostafavi, Hideyuki Yoshida, Devapregasan Moodley, Hugo LeBoité, Katherine Rothamel, Towfique Raj, Chun Jimmie Ye, Nicolas Chevrier, Shen-Ying Zhang, Ting Feng, et al. Parsing the interferon transcriptional network and its disease associations. Cell, 164(3):564–578, 2016.

[16] Ang Cui, Teddy Huang, Shuqiang Li, Aileen Ma, Jorge L Pérez, Chris Sander, Derin B Keskin, Catherine J Wu, Ernest Fraenkel, and Nir Hacohen. Dictionary of immune responses to cytokines at single-cell resolution. Nature, 625(7994):377–384, 2024.

[17] Seema S Ahuja, Fotini Paliogianni, Hidehiro Yamada, James E Balow, and DT Boumpas. Effect of transforming growth factor-beta on early and late activation events in human T cells. The Journal of Immunology, 150(8):3109– 3118, 1993.

[18] John J Bright, Lawrence D Kerr, and Subramaniam Sriram. TGF-beta inhibits IL-2-induced tyrosine phosphorylation and activation of Jak-1 and Stat 5 in T lymphocytes. The Journal of Immunology, 159(1):175–183, 1997.

[19] John J Bright and Subramaniam Sriram. TGF-β inhibits IL-12-induced activation of JAK-STAT pathway in T lymphocytes. The Journal of Immunology, 161(4):1772–1777, 1998.

[20] ćecile Pardoux, Xiaojing Ma, Stéphanie Gobert, Sandra Pellegrini, Patrick Mayeux, Françoise Gay, Giorgio Trinchieri, and Salem Chouaib. Downregulation of interleukin-12 (IL-12) responsiveness in human T cells by transforming growth factor-β:relationship with IL-12 signaling. Blood,The Journal of the American Society of Hematology, 93(5):1448–1455, 1999.

[21] Neha Cheemalavagu, Karsen E Shoger, Yuqi M Cao, Brandon A Michalides, Samuel A Botta, James R Faeder, and Rachel A Gottschalk. Predicting gene-level sensitivity to JAK-STAT signaling perturbation using a mechanistic-to-machine learning framework. Cell Systems, 15(1):37–48, 2024.

[22] Tadamitsu Kishimoto. Interleukin-6: discovery of a pleiotropic cytokine. Arthritis research & therapy, 8:1–6, 2006.

[23] Carly E Whyte, Kailash Singh, Oliver T Burton, Meryem Aloulou, Lubna Kouser, Rafael Valente Veiga, Amy Dashwood, Hanneke Okkenhaug, Samira Benadda, Alena Moudra, et al. Context-dependent effects of IL-2 rewire immunity into distinct cellular circuits. Journal of Experimental Medicine, 219(7):e20212391, 2022.

[24] Dominique Angèle Vuitton. The ambiguous role of immunity in echinococcosis: protection of the host or of the parasite? Acta tropica, 85(2):119–132, 2003.

[25] Francesco Annunziato, Chiara Romagnani, and Sergio Romagnani. The 3 major types of innate and adaptive cell-mediated effector immunity. Journal of Allergy and Clinical Immunology, 135(3):626–635, 2015.

[26] Vincenzo Barnaba, Marino Paroli, and Silvia Piconese. The ambiguity in immunology. Frontiers in immunology, 3:18, 2012.

[27] Cheol-Jung Lee, Hyun-Jung An, Eun Suh Cho, Han Chang Kang, Joo Young Lee, Hye Suk Lee, and Yong-Yeon Cho. Stat2 stability regulation: an intersection between immunity and carcinogenesis. Experimental & molecular medicine, 52(9):1526–1536, 2020.

[28] Ana L Romero-Weaver, Hsiang-Wen Wang, Håkan C Steen, Anthony J Scarzello, Veronica L Hall, Faruk Sheikh, Raymond P Donnelly, and Ana M Gamero. Resistance to IFN-α–induced apoptosis is linked to a loss of STAT2. Molecular Cancer Research, 8(1):80–92, 2010.

[29] Sanjay Gupta, Man Jiang, and Alessandra B Pernis. IFN-α activates stat6 and leads to the formation of stat2:stat6 complexes in B cells. The Journal of Immunology, 163(7):3834–3841, 1999.

[30] David M Frucht, Martin Aringer, Jérôme Galon, Carol Danning, Martin Brown, Samuel Fan, Michael Centola, Chang-You Wu, Nubuo Yamada, Hani El Gabalawy, et al. Stat4 is expressed in activated peripheral blood monocytes,dendritic cells,and macrophages at sites of Th1-mediated inflammation. The Journal of Immunology, 164(9):4659–4664, 2000.

[31] Tomohiro Yoshimoto, Kiyoshi Takeda, Takashi Tanaka, Kazunobu Ohkusu, Shin-ichiro Kashiwamura, Haruki Okamura, Shizuo Akira, and Kenji Nakanishi. IL-12 up-regulates IL-18 receptor expression on T cells, Th1 cells, and B cells:synergism with IL-18 for IFN-γ production. The Journal of Immunology, 161(7):3400–3407, 1998.

[32] Brian Orcutt-Jahns, Joao Rodrigues Lima Junior, Emily Lin, Russell C Rockne, Adina Matache, Sergio Branciamore, Ethan Hung, Andrei S Rodin, Peter P Lee, and Aaron S Meyer. Systems profiling reveals recurrently dysregulated cytokine signaling responses in ER+breast cancer patients’ blood. npj Systems Biology and Applications, 10(1):118, 2024.

[33] Francoise Cottrez and Hervé Groux. Regulation of TGF-β response during T cell activation is modulated by IL-10. The Journal of Immunology, 167(2):773–778, 2001.

[34] Caroline H Wallace, Bill X Wu, Mohammad Salem, Ephraim A Ansa-Addo, Alessandra Metelli, Shaoli Sun, Gary Gilkeson, Mark J Shlomchik, Bei Liu, and Zihai Li. B lymphocytes confer immune tolerance via cell surface GARP-TGF-β complex. JCI insight, 3(7), 2018.

[35] Aoife Kelly, Sezin Gunaltay, Craig P McEntee, Elinor E Shuttleworth, Catherine Smedley, Stephanie A Houston, Thomas M Fenton, Scott Levison, Elizabeth R Mann, and Mark A Travis. Human monocytes and macrophages regulate immune tolerance via integrin αvβ8-mediated TGF-β activation. Journal of Experimental Medicine, 215(11):2725–2736, 2018.

[36] Greg M Delgoffe and Dario AA Vignali. STAT heterodimers in immunity: A mixed message or a unique signal? Jak-stat, 2(1):e23060, 2013.

[37] Wei Liao, Jian-Xin Lin, and Warren J Leonard. IL-2 family cytokines:new insights into the complex roles of IL-2 as a broad regulator of T helper cell differentiation. Current opinion in immunology, 23(5):598–604, 2011.

[38] Kevin N Couper, Daniel G Blount, and Eleanor M Riley. IL-10:the master regulator of immunity to infection. The Journal of Immunology, 180(9):5771–5777, 2008.

[39] Tim R Mosmann. Regulation of immune responses by T cells with different cytokine secretion phenotypes: role of a new cytokine, cytokine synthesis inhibitory factor (il10). International Archives of Allergy and Immunology, 94(1-4):110–115, 1991.

[40] Kenji Itoh and Shunsei Hirohata. The role of IL-10 in human B cell activation, proliferation, and differentiation. The Journal of Immunology, 154(9):4341–4350, 1995.

[41] Catherine S Tripp, Stanley F Wolf, and Emil R Unanue. Interleukin 12 and tumor necrosis factor alpha are costimulators of interferon gamma production by natural killer cells in severe combined immunodeficiency mice with listeriosis, and interleukin 10 is a physiologic antagonist. Proceedings of the National Academy of Sciences, 90(8):3725–3729, 1993.

[42] Logan K Smith, Giselle M Boukhaled, Stephanie A Condotta, Sabrina Mazouz, Jenna J Guthmiller, Rahul Vijay, Noah S Butler, Julie Bruneau, Naglaa H Shoukry, Connie M Krawczyk, et al. Interleukin-10 directly inhibits CD8+ T cell function by enhancing N-glycan branching to decrease antigen sensitivity. Immunity, 48(2):299–312, 2018.

[43] Rene de Waal Malefyt, John Abrams, Bruce Bennett, Carl G Figdor, and Jan E de Vries. Interleukin 10 (IL-10) inhibits cytokine synthesis by human monocytes:an autoregulatory role of IL-10 produced by monocytes. The Journal of experimental medicine, 174(5):1209–1220, 1991.

[44] Kerstin Steinbrink, Matthias Wölfl, Helmut Jonuleit, Jurgen Knop, and Alexander H Enk. Induction of tolerance by IL-10-treated dendritic cells. The Journal of Immunology, 159(10):4772–4780, 1997.

[45] Valentina Carlini, Douglas M Noonan, Eslam Abdalalem, Delia Goletti, Clementina Sansone, Luana Calabrone, and Adriana Albini. The multifaceted nature of IL-10: regulation, role in immunological homeostasis and its relevance to cancer, COVID-19 and post-COVID conditions. Frontiers in immunology, 14:1161067, 2023.

[46] Yves Levy, Jean-Claude Brouet, et al. Interleukin-10 prevents spontaneous death of germinal center B cells by induction of the bcl-2 protein. The Journal of clinical investigation, 93(1):424–428, 1994.

[47] David F Fiorentino, Albert Zlotnik, Timothy R Mosmann, Maureen Howard, and Anne O’Garra. IL-10 inhibits cytokine production by activated macrophages. The Journal of Immunology, 147(11):3815–3822, 1991.

[48] Caroline Demangel, Patrick Bertolino, and Warwick J Britton. Autocrine IL-10 impairs dendritic cell (DC)-derived immune responses to mycobacterial infection by suppressing DC trafficking to draining lymph nodes and local IL-12 production. European journal of immunology, 32(4):994–1002, 2002.

[49] Lei Sun, Ren-Feng Guo, Michael W Newstead, Theodore J Standiford, Demetrio R Macariola, and Thomas P Shanley. Effect of IL-10 on neutrophil recruitment and survival after Pseudomonas aeruginosa challenge. American journal of respiratory cell and molecular biology, 41(1):76–84, 2009.

[50] Dajeong Lee, Min Geun Jo, Keun Young Min, Min Yeong Choi, Young Mi Kim, Hyuk Soon Kim, and Wahn Soo Choi. IL-10+ regulatory B cells mitigate atopic dermatitis by suppressing eosinophil activation. Scientific Reports, 14(1):18164, 2024.

[51] Shaoxuan Wang, JinXuan Wang, Ran Ma, Shaofeng Yang, Tingting Fan, Jing Cao, Yang Wang, Wenbin Ma, Wenxiu Yang, Fulai Wang, et al. IL-10 enhances T cell survival and is associated with faster relapse in patients with inactive ulcerative colitis. Molecular immunology, 121:92–98, 2020.

[52] Eric Lelievre, Denis Sarrouilhe, Franck Morel, Jean-Louis Preud’Homme, John Wijdenes, and Jean-Claude Lecron. Preincubation of human resting T cell clones with interleukin 10 strongly enhances their ability to produce cytokines after stimulation. Cytokine, 10(11):831–840, 1998.

[53] Francoise Rousset, Eric Garcia, Thierry Defrance, Catherine Peronne, Nadia Veïo, Di-Hwei Hsu, Rob Kastelein, Kevin W Moore, and Jacques Banchereau. Interleukin 10 is a potent growth and differentiation factor for activated human B lymphocytes. Proceedings of the National Academy of Sciences, 89(5):1890–1893, 1992.

[54] Zixi Wang, D. Guan, Jianxin Huo, Subhra K Biswas, Yuhan Huang, Yuansheng Yang, Shengli Xu, and Kong-Peng Lam. IL-10 enhances human natural killer cell effector functions via metabolic reprogramming regulated by mtorc1 signaling. Frontiers in immunology, 12:619195, 2021.

[55] Simone Mocellin, M Panelli, E Wang, Carlo Riccardo Rossi, Pierluigi Pilati, Donato Nitti, Mario Lise, and FM Marincola. IL-10 stimulatory effects on human NK cells explored by gene profile analysis. Genes & Immunity, 5(8):621–630, 2004.

[56] Martin A Schwarz, Linda D Hamilton, Lisa Tardelli, Satwant K Narula, and Lee M Sullivan. Stimulation of cytolytic activity by interleukin-10. Journal of immunotherapy, 16(2):95–104, 1994.

[57] Kazunaga Agematsu, Haruo Nagumo, Yumiko Oguchi, Takayuki Nakazawa, Keitaro Fukushima, Kozo Yasui, Susumu Ito, Tetsuji Kobata, Chikao Morimoto, and Atsushi Komiyama. Generation of plasma cells from peripheral blood memory B cells: synergistic effect of interleukin-10 and CD27/CD70 interaction. Blood,The Journal of the American Society of Hematology, 91(1):173–180, 1998.

[58] Jan Emmerich, John B Mumm, Ivan H Chan, Drake LaFace, Hoa Truong, Terrill McClanahan, Daniel M Gorman, and Martin Oft. IL-10 directly activates and expands tumor-resident CD8+ T cells without de novo infiltration from secondary lymphoid organs. Cancer research, 72(14):3570–3581, 2012.

[59] John B Mumm, Jan Emmerich, Xueqing Zhang, Ivan Chan, Lingling Wu, Smita Mauze, Steven Blaisdell, Beth Basham, Jie Dai, Jeff Grein, et al. IL-10 elicits IFNγ-dependent tumor immune surveillance. Cancer cell, 20(6):781–796, 2011.

[60] Shin-ichiro Fujii, Kanako Shimizu, Takashi Shimizu, and Michael T Lotze. Interleukin-10 promotes the maintenance of antitumor CD8+ T-cell effector function in situ. Blood,The Journal of the American Society of Hematology, 98(7):2143–2151, 2001.

[61] Bruce R Blazar, Patricia A Taylor, Angela Panoskaltsis-Mortari, Satwant K Narula, Sidney R Smith, Maria G Roncarolo, and Daniel A Vallera. Interleukin-10 dose-dependent regulation of CD4+and CD8+T cell-mediated graft-versus-host disease. Transplantation, 66(9):1220–1229, 1998.

[62] Li-Mou Zheng, David M Ojcius, F Garaud, C Roth, E Maxwell, Z Li, H Rong, J Chen, XY Wang, JJ Catino, et al. Interleukin-10 inhibits tumor metastasis through an NK cell-dependent mechanism. The Journal of experimental medicine, 184(2):579–584, 1996.

[63] Robert M Berman, Tadamichi Suzuki, Hideaki Tahara, Paul D Robbins, Satwant K Narula, and Michael T Lotze. Systemic administration of cellular IL-10 induces an effective, specific, and long-lived immune response against established tumors in mice. The Journal of Immunology, 157(1):231–238, 1996.

[64] H Tilg, C Van Montfrans, A Van den Ende, A Kaser, SJH Van Deventer, S Schreiber, M Gregor, O Ludwiczek, P Rutgeerts, C Gasche, et al. Treatment of Crohn’s disease with recombinant human interleukin 10 induces the proinflammatory cytokine interferon-γ. Gut, 50(2):191–195, 2002.

[65] Richard N Fedorak, Alfred Gangl, Charles O Elson, Paul Rutgeerts, Stefan Schreiber, Gary Wild, Stephen B Hanauer, Ann Kilian, Marielle Cohard, Alexandre LeBeaut, et al. Recombinant human interleukin 10 in the treatment of patients with mild to moderately active Crohn’s disease. Gastroenterology, 119(6):1473–1482, 2000.

[66] Aung Naing, Jeffrey R Infante, Kyriakos P Papadopoulos, Ivan H Chan, Cong Shen, Navneet P Ratti, Bianca Rojo, Karen A Autio, Deborah J Wong, Manish R Patel, et al. PEGylated IL-10 (Pegilodecakin) induces systemic immune activation, CD8+ T cell invigoration and polyclonal T cell expansion in cancer patients. Cancer Cell, 34(5):775–791, 2018.

[67] Fanny N Lauw, Dasja Pajkrt, C Erik Hack, Masashi Kurimoto, Sander JH van Deventer, and Tom van der Poll. Proinflammatory effects of IL-10 during human endotoxemia. The Journal of Immunology, 165(5):2783–2789, 2000.

[68] Richard D Huhn, Elaine Radwanski, Sean M O’Connell, Marc G Sturgill, Laura Clarke, Ronald P Cody, Melton B Affrime, and David L Cutler. Pharmacokinetics and immunomodulatory properties of intravenously administered recombinant human interleukin-10 in healthy volunteers. Blood, 87(2):699–705, 1996.

[69] Huan Han, Qingfeng Ma, Cong Li, Rui Liu, Li Zhao, Wei Wang, Pingan Zhang, Xinghui Liu, Guosheng Gao, Fang Liu, et al. Profiling serum cytokines in COVID-19 patients reveals IL-6 and IL-10 are disease severity predictors. Emerging microbes & infections, 9(1):1123–1130, 2020.

[70] Zhe Xu, Lei Shi, Yijin Wang, Jiyuan Zhang, Lei Huang, Chao Zhang, Shuhong Liu, Peng Zhao, Hongxia Liu, Li Zhu, et al. Pathological findings of COVID-19 associated with acute respiratory distress syndrome. The Lancet respiratory medicine, 8(4):420–422, 2020.

[71] Ligong Lu, Hui Zhang, Danielle J Dauphars, and You-Wen He. A potential role of interleukin 10 in COVID-19 pathogenesis. Trends in Immunology, 42(1):3–5, 2021.

[72] Hashim Islam, Thomas C Chamberlain, Alice L Mui, and Jonathan P Little. Elevated interleukin-10 levels in COVID-19: potentiation of pro-inflammatory responses or impaired anti-inflammatory action? Frontiers in Immunology, 12:677008, 2021.

[73] Nicole Prince, Julia A Penatzer, Taylor L Shackleford, Elizabeth K Stewart, Matthew J Dietz, and Jonathan W Boyd. Tissue-level cytokines in a rodent model of chronic implant-associated infection. Journal of Orthopaedic Research®, 39(10):2159–2168, 2021.

[74] Huixun Du, Juliet M Bartleson, Sergei Butenko, Valentina Alonso, Wendy F Liu, Daniel A Winer, and Manish J Butte. Tuning immunity through tissue mechanotransduction. Nature Reviews Immunology, 23(3):174–188, 2023.

[75] Lital Mordechay, Guillaume Le Saux, Avishay Edri, Uzi Hadad, Angel Porgador, and Mark Schvartzman. Mechanical regulation of the cytotoxic activity of natural killer cells. ACS Biomaterials Science & Engineering, 7(1):122–132, 2020.

[76] Hamza Atcha, Amit Jairaman, Jesse R Holt, Vijaykumar S Meli, Raji R Nagalla, Praveen Krishna Veerasubramanian, Kyle T Brumm, Huy E Lim, Shivashankar Othy, Michael D Cahalan, et al. Mechanically activated ion channel Piezo1 modulates macrophage polarization and stiffness sensing. Nature communications, 12(1):3256, 2021.

[77] Samina Shaheen, Zhengpeng Wan, Zongyu Li, Alicia Chau, Xinxin Li, Shaosen Zhang, Yang Liu, Junyang Yi, Yingyue Zeng, Jing Wang, et al. Substrate stiffness governs the initiation of B cell activation by the concerted signaling of PKCβ and focal adhesion kinase. Elife, 6:e23060, 2017.

[78] Michael Saitakis, Stéphanie Dogniaux, Christel Goudot, Nathalie Bufi, Sophie Asnacios, Mathieu Maurin, Clotilde Randriamampita, Atef Asnacios, and Claire Hivroz. Different TCR-induced T lymphocyte responses are potentiated by stiffness with variable sensitivity. Elife, 6:e23190, 2017.

[79] Kevin P Meng, Fatemeh S Majedi, Timothy J Thauland, and Manish J Butte. Mechanosensing through YAP controls T cell activation and metabolism. Journal of Experimental Medicine, 217(8):e20200053, 2020.

[80] Bola S Hanna, Laura Llao-Cid, Murat Iskar, Philipp M Roessner, Lara C Klett, John KL Wong, Yashna Paul, Nikolaos Ioannou, Selcen Öztürk, Norman Mack, et al. Interleukin-10 receptor signaling promotes the maintenance of a PD-1int TCF-1+CD8+ T cell population that sustains anti-tumor immunity. Immunity, 54(12):2825–2841, 2021.

[81] Nir Yogev, Tanja Bedke, Yasushi Kobayashi, Leonie Brockmann, Dominika Lukas, Tommy Regen, Andrew L Croxford, Alexei Nikolav, Nadine Hövelmeyer, Esther von Stebut, et al. CD4+ T-cell-derived IL-10 promotes CNS inflammation in mice by sustaining effector T cell survival. Cell Reports, 38(13), 2022.

[82] Margarida Saraiva, Paulo Vieira, and Anne O’garra. Biology and therapeutic potential of interleukin-10. Journal of Experimental Medicine, 217(1):e20190418, 2019.

[83] Toshio Tanaka, Masashi Narazaki, and Tadamitsu Kishimoto. IL-6 in inflammation, immunity, and disease. Cold Spring Harbor perspectives in biology, 6(10):a016295, 2014.

[84] Sean Diehl, Chi-Wing Chow, Linda Weiss, Alois Palmetshofer, Thomas Twardzik, Laura Rounds, Edgar Serfling, Roger J Davis, Juan Anguita, and Mercedes Rinćon. Induction of NFATc2 expression by interleukin 6 promotes T helper type 2 differentiation. The Journal of experimental medicine, 196(1):39–49, 2002.

[85] Nicola M Heller, William M Gwinn, Raymond P Donnelly, Stephanie L Constant, and Achsah D Keegan. IL-4 engagement of the type I IL-4 receptor complex enhances mouse eosinophil migration to eotaxin-1 in vitro. PLoS One, 7(6):e39673, 2012.

[86] Monica I Vazquez, Jovani Catalan-Dibene, and Albert Zlotnik. B cells responses and cytokine production are regulated by their immune microenvironment. Cytokine, 74(2):318–326, 2015.

[87] Olof Berggren, Andrei Alexsson, David L Morris, Karolina Tandre, Gert Weber, Timothy J Vyse, Ann-Christine Syvänen, Lars Rönnblom, and Maija-Leena Eloranta. IFN-α production by plasmacytoid dendritic cell associations with polymorphisms in gene loci related to autoimmune and inflammatory diseases. Human Molecular Genetics, 24(12):3571–3581, 2015.

[88] Stefania Gallucci, Martijn Lolkema, and Polly Matzinger. Natural adjuvants: endogenous activators of dendritic cells. Nature medicine, 5(11):1249–1255, 1999.

[89] Jennifer Martinez, Xiaopei Huang, and Yiping Yang. Direct action of type I IFN on NK cells is required for their activation in response to vaccinia viral infection in vivo. The Journal of Immunology, 180(3):1592–1597, 2008.

[90] Alsya J Affandi, Katarzyna Olesek, Joanna Grabowska, Maarten K Nijen Twilhaar, Ernesto Rodríguez, Anno Saris, Eline S Zwart, Esther J Nossent, Hakan Kalay, Michael De Kok, et al. CD169 defines activated CD14+ monocytes with enhanced CD8+ T cell activation capacity. Frontiers in immunology, 12:697840, 2021.

[91] Déborah Braun, Iris Caramalho, and Jocelyne Demengeot. IFN-α/β enhances BCR-dependent B cell responses. International immunology, 14(4):411–419, 2002.

[92] Julie M Curtsinger, Javier O Valenzuela, Pujya Agarwal, Debra Lins, and Matthew F Mescher. Cutting edge: type I IFNs provide a third signal to CD8 T cells to stimulate clonal expansion and differentiation. The Journal of Immunology, 174(8):4465–4469, 2005.

[93] Jianping Pan, Minghui Zhang, Jianli Wang, Qingqing Wang, Dajing Xia, Wenji Sun, Lihuang Zhang, Hai Yu, Yongjun Liu, and Xuetao Cao. Interferon-γ is an autocrine mediator for dendritic cell maturation. Immunology letters, 94(1-2):141–151, 2004.

[94] Purnima Bhat, Graham Leggatt, Nigel Waterhouse, and Ian H Frazer. Interferon-γ derived from cytotoxic lymphocytes directly enhances their motility and cytotoxicity. Cell death & disease, 8(6):e2836–e2836, 2017.

[95] Rosario Luque-Martin, Davina C Angell, Mathias Kalxdorf, Sharon Bernard, William Thompson, H Christian Eberl, Charlotte Ashby, Johannes Freudenberg, Catriona Sharp, Jan Van den Bossche, et al. IFN-γ drives human monocyte differentiation into highly proinflammatory macrophages that resemble a phenotype relevant to psoriasis. The Journal of Immunology, 207(2):555–568, 2021.

[96] Cristiane Jaciara Furlaneto and Ana Campa. A novel function of serum amyloid A: a potent stimulus for the release of tumor necrosis factor-α, interleukin-1β,and interleukin-8 by human blood neutrophil. Biochemical and biophysical research communications, 268(2):405–408, 2000.

[97] Priscilla Biswas, Fanny Delfanti, Sergio Bernasconi, Manuela Mengoï, Manuela Cota, Nadia Polentarutti, Alberto Mantovani, Adriano Laärin, Silvano Soäni, and Guido Poli. Interleukin-6 induces monocyte chemotactic protein-1 in peripheral blood mononuclear cells and in the U937 cell line. Blood,The Journal of the American Society of Hematology, 91(1):258–265, 1998.

[98] Jamie R Schoenborn and Christopher B Wilson. Regulation of interferon-γ during innate and adaptive immune responses. Advances in immunology, 96:41–101, 2007.

[99] P Sachin, S Gadani, J Cronk, G Norris, and J Kipnis. Interleukin-4: a cytokine to remember. The Journal of Immunology, 189:4213–4421, 2012.

[100] Li-Rung Huang, Fen-Ling Chen, Yi-Ting Chen, Ya-Min Lin, and John T Kung. Potent induction of long-term CD8+ T cell memory by short-term IL-4 exposure during t cell receptor stimulation. Proceedings of the National Academy of Sciences, 97(7):3406–3411, 2000.

[101] Volker Brinkmann, Thomas Geiger, Sefik Alkan, and Christoph H Heusser. Interferon alpha increases the frequency of interferon gamma-producing human CD4+ T cells. The Journal of experimental medicine, 178(5):1655–1663, 1993.

[102] Jennifer M Reed, Patrick J Branigan, and Anil Bamezai. Interferon gamma enhances clonal expansion and survival of CD4+ T cells. Journal of interferon & cytokine research, 28(10):611–622, 2008.

[103] Arianexys Aquino-López, Vladimir V Senyukov, Zlatko Vlasic, Eugenie S Kleinerman, and Dean A Lee. Interferon gamma induces changes in natural killer (NK) cell ligand expression and alters NK cell-mediated lysis of pediatric cancer cell lines. Frontiers in immunology, 8:391, 2017.

[104] Arnon Nagler, Lewis L Lanier, and Joseph H Phillips. The effects of IL-4 on human natural killer cells. A potent regulator of il-2 activation and proliferation. The Journal of Immunology, 141(7):2349–2351, 1988.

[105] Richard Essner, Kristina Rhoades, William H McBride, Donald L Morton, and JS Economou. IL-4 down-regulates IL-1 and TNF gene expression in human monocytes. The Journal of Immunology, 142(11):3857–3861, 1989.

[106] Ryotaro Yoshida, Henry W Murray, and Carl F Nathan. Agonist and antagonist effects of interferon alpha and beta on activation of human macrophages. Two classes of interferon gamma receptors and blockade of the high-affinity sites by interferon alpha or beta. The Journal of experimental medicine, 167(3):1171–1185, 1988.

[107] Uma Sriram, Chhanda Biswas, Edward M Behrens, Joudy-Ann Dinnall, Debra K Shivers, Marc Monestier, Yair Argon, and Stefania Gallucci. IL-4 suppresses dendritic cell response to type I interferons. The Journal of Immunology, 179(10):6446–6455, 2007.

[108] Maili Zimmermann, Fabio Arruda-Silva, Francisco Bianchetto-Aguilera, Giulia Finotti, Federica Calzetti, Patrizia Scapini, Claudio Lunardi, Marco A Cassatella, and Nicola Tamassia. IFNα enhances the production of IL-6 by human neutrophils activated via TLR8. Scientific reports, 6(1):19674, 2016.

[109] Sibylla Martinelli, Mirjana Urosevic, Arezoo Daryadel, Patrick Antony Oberholzer, Christa Baumann, Martin F Fey, Reinhard Dummer, Hans-Uwe Simon, and Shida Yousefi. Induction of genes mediating interferon-dependent extracellular trap formation during neutrophil differentiation. Journal of Biological Chemistry, 279(42):44123–44132, 2004.

[110] Paige Lacy, Francesca Levi-Schaffer, Salahaddin Mahmudi-Azer, Ben Bablitz, Stacey C Hagen, Juan Velazquez, A Barry Kay, and Redwan Moqbel. Intracellular localization of interleukin-6 in eosinophils from atopic asthmatics and effects of interferon-γ. Blood,The Journal of the American Society of Hematology, 91(7):2508–2516, 1998.

[111] Delphine Aldebert, Bouchäib Lamkhioued, Corinne Desaint, Abdelilah Soussi Gounni, Michel Goldman, André Capron, Lionel Prin, and Monique Capron. Eosinophils express a functional receptor for interferon alpha: inhibitory role of interferon alpha on the release of mediators. Blood, 87(6):2354–2360, 1996.

[112] Akira Kanda, Virginie Driss, Nicolas Hornez, Marwan Abdallah, Thomas Roumier, Georges Abboud, Fanny Legrand, Delphine Staumont-Sallé, Severine Quéant, Julie Bertout, et al. Eosinophil-derived IFN-γ induces airway hyperresponsiveness and lung inflammation in the absence of lymphocytes. Journal of allergy and clinical immunology, 124(3):573–582, 2009.

[113] Melissa Swiecki and Marco Colonna. Type I interferons: diversity of sources, production pathways and effects on immune responses. Current opinion in virology, 1(6):463–475, 2011.

[114] Karsten Wessel Eriksen, Viveca Horst Sommer, Anders Woetmann, Anette Bødker Rasmussen, Christine Brender, Arne Svejgaard, Søren Skov, Carsten Geisler, and Niels Ødum. Bi-phasic effect of interferon (IFN)-α: IFN-α up-and down-regulates interleukin-4 signaling in human T cells. Journal of Biological Chemistry, 279(1):169–176, 2004.

[115] Janine Woytschak, Nadia Keller, Carsten Krieg, Daniela Impelliïeri, Robert W Thompson, Thomas A Wynn, Annelies S Zinkernagel, and Onur Boyman. Type 2 interleukin-4 receptor signaling in neutrophils antagonizes their expansion and migration during infection and inflammation. Immunity, 45(1):172–184, 2016.

[116] Cong Wu, Yiquan Xue, Pin Wang, Li Lin, Qiuyan Liu, Nan Li, Junfang Xu, and Xuetao Cao. IFN-γ primes macrophage activation by increasing phosphatase and tensin homolog via downregulation of miR-3473b. The Journal of Immunology, 193(6):3036–3044, 2014.

[117] Siamon Gordon and Fernando O Martinez. Alternative activation of macrophages: mechanism and functions. Immunity, 32(5):593–604, 2010.

[118] Jan Ackermann, Lilli Arndt, Janine Fröba, Andreas Lindhorst, Markus Glaß, Michaela Kirstein, Constance Hobusch, F Thomas Wunderlich, Julia Braune, and Martin Gericke. IL-6 signaling drives self-renewal and alternative activation of adipose tissue macrophages. Frontiers in Immunology, 15:1201439, 2024.

[119] Song Guo Zheng, Juhua Wang, and David A Horwitz. Cutting edge: Foxp3+CD4+CD25+ regulatory T cells induced by IL-2 and TGF-β are resistant to Th17 conversion by IL-6. The Journal of Immunology, 180(11):7112–7116, 2008.

[120] Ronald B Smeltz, June Chen, Rolf Ehrhardt, and Ethan M Shevach. Role of IFN-γ in Th1 differentiation: IFN-γ regulates IL-18Rα expression by preventing the negative effects of IL-4 and by inducing/maintaining IL-12 receptor β2 expression. The Journal of Immunology, 168(12):6165–6172, 2002.

[121] Susan L Swain, Andrew D Weinberg, Michele English, and Gail Huston. IL-4 directs the development of Th2-like helper effectors. The Journal of Immunology, 145(11):3796–3806, 1990.

[122] Loredana Cifaldi, Giusi Prencipe, Ivan Caiello, Claudia Bracaglia, Franco Locatelli, Fabrizio De Benedetti, and Raffaele Strippoli. Inhibition of natural killer cell cytotoxicity by interleukin-6: implications for the pathogenesis of macrophage activation syndrome. Arthritis & rheumatology, 67(11):3037–3046, 2015.

[123] Sung-Joo Park, Takayuki Nakagawa, Hidemitsu Kitamura, Toru Atsumi, Hokuto Kamon, Shin-ichiro Sawa, Daisuke Kamimura, Naoko Ueda, Yoichiro Iwakura, Katsuhiko Ishihara, et al. IL-6 regulates in vivo dendritic cell differentiation through STAT3 activation. The Journal of Immunology, 173(6):3844–3854, 2004.

[124] Takayuki Nakagawa, Mineko Tsuruoka, Hideki Ogura, Yuko Okuyama, Yasunobu Arima, Toshio Hirano, and Masaaki Murakami. IL-6 positively regulates Foxp3+CD8+ T cells in vivo. International immunology, 22(2):129–139, 2010.

[125] Rui Yang, April R Masters, Karen A Fortner, Devin P Champagne, Natalia Yanguas-Casás, Daniel J Silberger, Casey T Weaver, Laura Haynes, and Mercedes Rincon. IL-6 promotes the differentiation of a subset of naive CD8+T cells into IL-21-producing B helper CD8+ T cells. Journal of Experimental Medicine, 213(11):2281–2291, 2016.

[126] Nicholas E Martinez, Fumitaka Sato, Eiichiro Kawai, Seiichi Omura, Robert P Chervenak, and Ikuo Tsunoda. Regulatory T cells and Th17 cells in viral infections: implications for multiple sclerosis and myocarditis. Future virology, 7(6):593–608, 2012.

[127] Dan Aderka, JM Le, and J Vilcek. IL-6 inhibits lipopolysaccharide-induced tumor necrosis factor production in cultured human monocytes, U937 cells, and in mice. The Journal of Immunology, 143(11):3517–3523, 1989.

[128] Weimin Wu, Kirsten K Dietze, Kathrin Gibbert, Karl S Lang, Mirko Trilling, Huimin Yan, Jun Wu, Dongliang Yang, Mengji Lu, Michael Roggendorf, et al. TLR ligand induced IL-6 counter-regulates the anti-viral CD8+T cell response during an acute retrovirus infection. Scientific reports, 5(1):10501, 2015.

[129] Sean B Joseph, Kent T Miner, and Michael Croft. Augmentation of naive, Th1 and Th2 effector CD4 responses by IL-6, IL-1 and TNF. European journal of immunology, 28(1):277–289, 1998.

[130] Suzanne C Morris, Tatyana Orekhova, Michelle J Meadows, Stephanie M Heidorn, Junqi Yang, and Fred D Finkelman. IL-4 induces in vivo production of IFN-γ by NK and NKT cells. The Journal of Immunology, 176(9):5299–5305, 2006.

[131] H Rabinowich, P Sedlmayr, RB Herberman, and TL Whiteside. Response of human NK cells to IL-6 alterations of the cell surface phenotype,adhesion to fibronectin and laminin,and tumor necrosis factor-alpha/beta secretion. The Journal of Immunology, 150(11):4844–4855, 1993.

[132] Tzer-Min Kuo, Cheng-po Hu, Ya-Ling Chen, Ming-Hsiang Hong, King-Song Jeng, Chun-Chin T Liang, Mong-Liang Chen, and Chungming Chang. HBV replication is significantly reduced by IL-6. Journal of biomedical science, 16:1–9, 2009.

[133] Keila Zaniboni Siqueira, Ângela Maria Victoriano De Campos Soares, Luciane Alarcão Dias-Melicio, Sueli Aparecida Calvi, and Maria Terezinha Serrão Peraçoli. Interleukin-6 treatment enhances human monocyte permissiveness for Paracoccidioides brasiliensis growth by modulating cytokine production. Medical Mycology, 47(3):259–267, 2009.

[134] Teresa Zelante, Jan Fric, Alicia YW Wong, and Paola Ricciardi-Castagnoli. Interleukin-2 production by dendritic cells and its immuno-regulatory functions. Frontiers in immunology, 3:161, 2012.

[135] Denis Girard, Jean Gosselin, Dominique Heitz, Robert Paquin, and Andrε D Beaulieu. Effects of interleukin-2 on gene expression in human neutrophils. Blood, 86:1170—1176, 1995.

[136] TH Rand, DS Silberstein, H Kornfeld, PF Weller, et al. Human eosinophils express functional interleukin-2 receptors. The Journal of clinical investigation, 88(3):825–832, 1991.

[137] Ronald D Curran, Timothy R Billiar, Michael A West, Brandon G Bentz, and Richard L Simmons. Effect of interleukin-2 on kupffer cell activation: Interleukin-2 primes and activates Kupffer cells to suppress hepatocyte protein synthesis in vitro. Archives of Surgery, 123(11):1373–1378, 1988.

[138] Onur Boyman and Jonathan Sprent. The role of interleukin-2 during homeostasis and activation of the immune system. Nature Reviews Immunology, 12(3):180–190, 2012.

[139] Jean Dunne, Sara Lynch, Cliona O’Farrelly, Stephen Todryk, John E Hegarty, Conleth Feighery, and Derek G Doherty. Selective expansion and partial activation of human NK cells and NK receptor-positive T cells by IL-2 and IL-15. The Journal of Immunology, 167(6):3129–3138, 2001.

[140] Nayoung Kim, Eunbi Yi, Eunbi Lee, Hyo Jin Park, and Hun Sik Kim. Interleukin-2 is required for NKp30-dependent NK cell cytotoxicity by preferentially regulating NKp30 expression. Frontiers in Immunology, 15:1388018, 2024.

[141] Atsushi Muraguchi, John H Kehrl, Dan L Longo, David J Volkman, KendallA Smith, and Anthony S Fauci. Interleukin 2 receptors on human B cells. Implications for the role of interleukin-2 in human B cell function. The Journal of experimental medicine, 161(1):181–197, 1985.

[142] John Yates, Flavia Rovis, Peter Mitchell, Behdad Afzali, JY-S Tsang, Marina Garin, RI Lechler, Giovanna Lombardi, and OA Garden. The maintenance of human CD4+CD25+ regulatory T cell function: IL-2, IL-4, IL-7 and IL-15 preserve optimal suppressive potency in vitro. International immunology, 19(6):785–799, 2007.

[143] Maria C Bosco, Rafael E Curiel, Arnold H Zea, Maria G Malabarba, John R Ortaldo, and Igor Espinoza-Delgado. IL-2 signaling in human monocytes involves the phosphorylation and activation of p59hck 1. The Journal of Immunology, 164(9):4575–4585, 2000.

[144] M Misago, J Tsukada, R Ogawa, M Kikuchi, T Hanamura, S Chiba, S Oda, I Morimoto, and S Eto. Enhancing effects of IL-2 on M-CSF production by human peripheral blood monocytes. International Journal of Hematology, 58(1-2):43–51, 1993.

[145] MA Brach, C Arnold, M Kiehntopf, HJ Gruss, and F Herrmann. Transcriptional activation of the macrophage colony-stimulating factor gene by IL-2 is associated with secretion of bioactive macrophage colony-stimulating factor protein by monocytes and involves activation of the transcription factor NF-kappaB. Journal of immunology (Baltimore,Md.:1950), 150(12):5535–5543, 1993.

[146] I Espinoza-Delgado, DL Longo, GL Gusella, and L Varesio. IL-2 enhances c-fms expression in human monocytes. Journal of immunology (Baltimore,Md.:1950), 145(4):1137–1143, 1990.

[147] L Steven. Interleukin-2-induced tumor necrosis factor-alpha (TNF-gene expression in human alveolar macrophages and blood monocytes1-3. The Journal of Immunology, 151:2725–2732, 1993.

[148] GL Gusella, Tiziana Musso, MC Bosco, I Espinoza-Delgado, K Matsushima, and L Varesio. IL-2 up-regulates but IFN-gamma suppresses IL-8 expression in human monocytes. Journal of immunology (Baltimore,Md.:1950), 151(5):2725–2732, 1993.

[149] J Stankova, G Dupuis, N Gagnon, M Thivierge, S Turcotte, and M Rola-Pleszczynski. Priming of human monocytes with leukotriene B4 enhances their sensitivity in IL-2-driven tumor necrosis factor-alpha production. Transcriptional and post-transcriptional up-regulation of IL-2 receptors. Journal of immunology (Baltimore,Md.:1950), 150(9):4041–4051, 1993.

[150] PK Epling-Burnette, Sheng Wei, DK Blanchard, Emma Spranzi, and Julie Y Djeu. Coinduction of granulocyte-macrophage colony-stimulating factor release and lymphokine-activated killer cell susceptibility in monocytes by interleukin-2 via interleukin-2 receptor beta. Blood, 81(11):3130–3137, 1993.

[151] ML Lohmann-Matthes, A Emmendoerffer, and Li Hao. Influence of interleukin-2 on the differentiation of macrophages. Pathobiology, 59(3):117–121, 1991.

[152] Guan-Feng Ouyang, Masanao Saio, Tatsuhiko Suwa, Hisashi Imai, Jiro Nakagawa, Kenichi Nonaka, Naoki Umemura, Mika Kijima, and Tsuyoshi Takami. Interleukin-2 augmented activation of tumor associated macrophage plays the main role in MHC class I in vivo induction in tumor cells that are MHC negative in vitro. International journal of oncology, 28(5):1201–1208, 2006.

[153] Thomas R Malek and Iris Castro. Interleukin-2 receptor signaling: at the interface between tolerance and immunity. Immunity, 33(2):153–165, 2010.

[154] Masao Hashimoto, Se Jin Im, Koichi Araki, and Rafi Ahmed. Cytokine-mediated regulation of CD8 T-cell responses during acute and chronic viral infection. Cold Spring Harbor perspectives in biology, 11(1):a028464, 2019.

[155] ARNON Nagler, LEWIS L Lanier, and Joseph H Phillips. Constitutive expression of high affinity interleukin-2 receptors on human CD16-natural killer cells in vivo. The Journal of experimental medicine, 171(5):1527–1533, 1990.

[156] Akimichi Inaba, Zewen Kelvin Tuong, Tian X Zhao, Andrew P Stewart, Rebeccah Mathews, Lucy Truman, Rouchelle Sriranjan, Jane Kennet, Kourosh Saeb-Parsy, Linda Wicker, et al. Low-dose IL-2 enhances the generation of IL-10-producing immunoregulatory B cells. Nature Communications, 14(1):2071, 2023.

[157] Hyoung Pyo Kim, Jean Imbert, and Warren J Leonard. Both integrated and differential regulation of components of the IL-2/IL-2 receptor system. Cytokine & growth factor reviews, 17(5):349–366, 2006.

[158] Julie M Mazet, Jagdish N Mahale, Orion Tong, Robert A Watson, Ana Victoria Lechuga-Vieco, Gabriela Pirgova, Vivian WC Lau, Moustafa Attar, Lada A Koneva, Stephen N Sansom, et al. IFNγ signaling in cytotoxic T cells restricts anti-tumor responses by inhibiting the maintenance and diversity of intra-tumoral stem-like T cells. Nature communications, 14(1):321, 2023.

[159] Ryan S Waters, Justin SA Perry, SunPil Han, Bibiana Bielekova, and Tomas Gedeon. The effects of interleukin-2 on immune response regulation. Mathematical Medicine and Biology:a Journal of the IMA, 35(1):79–119, 2018.

[160] Michael Osherov and Ron Milo. Daclizumab for the treatment of adults with relapsing forms of multiple sclerosis. Expert Review of Clinical Pharmacology, 10(10):1037–1047, 2017.

[161] Takuya Yamaguchi, Chia Jung Chang, Axel Karger, Markus Keller, Florian Pfaff, Eakapol Wangkahart, Tiehui Wang, Christopher J Secombes, Azusa Kimoto, Mitsuru Furihata, et al. Ancient cytokine interleukin 15-like (IL-15L) induces a type 2 immune response. Frontiers in Immunology, 11:549319, 2020.

[162] Susana Pesoa, Andrea Martin, Ana Lia Mariani, Carlos Vullo, and Horacio Serra. Interleukin 2 induction of proliferation in resting T lymphocytes requires contact with monocytes. Medicina-Buenos Aires-, 60(2):202–210, 2000.

[163] Jens HW Pahl, Adelheid Cerwenka, and Jing Ni. Memory-like NK cells: remembering a previous activation by cytokines and NK cell receptors. Frontiers in immunology, 9:2796, 2018.

[164] Azucena Salas, Cristian Hernandez-Rocha, Marjolijn Duijvestein, William Faubion, Dermot McGovern, Severine Vermeire, Stefania Vetrano, and Niels Vande Casteele. JAK–STAT pathway targeting for the treatment of inflammatory bowel disease. Nature reviews Gastroenterology & hepatology, 17(6):323–337, 2020.

[165] Juan L Mendoza, Nichole K Escalante, Kevin M Jude, Junel Sotolongo Bellon, Leon Su, Tim M Horton, Naotaka Tsutsumi, Steven J Berardinelli, Robert S Haltiwanger, Jacob Piehler, et al. Structure of the IFNγ receptor complex guides design of biased agonists. Nature, 567(7746):56–60, 2019.

[166] Robert A Saxton, Naotaka Tsutsumi, Leon L Su, Gita C Abhiraman, Kritika Mohan, Lukas T Henneberg, Nanda G Aduri, Cornelius Gati, and K Christopher Garcia. Structure-based decoupling of the pro-and anti-inflammatory functions of interleukin-10. Science, 371(6535):eabc8433, 2021.

[167] Georgios Skiniotis, Martin J Boulanger, K Christopher Garcia, and Thomas Walz. Signaling conformations of the tall cytokine receptor gp130 when in complex with IL-6 and IL-6 receptor. Nature structural & molecular biology, 12(6):545–551, 2005.

[168] Hongchun Li, Nanaocha Sharma, Ignacio J General, Gideon Schreiber, and Ivet Bahar. Dynamic modulation of binding affinity as a mechanism for regulating interferon signaling. Journal of molecular biology, 429(16):2571–2589, 2017.

[169] Peter Holicek, Emma Guilbaud, Vanessa Klapp, Iva Truxova, Radek Spisek, Lorenzo Gallüi, and Jitka Fucikova. Type I interferon and cancer. Immunological Reviews, 321(1):115–127, 2024.

[170] Vanessa S Cull, Peta A Tilbrook, Emmalene J Bartlett, Natalie L Brekalo, and Cassandra M James. Type I interferon differential therapy for erythroleukemia: specificity of STAT activation. Blood,The Journal of the American Society of Hematology, 101(7):2727–2735, 2003.

[171] Bernardetta Nardelli, Liubov Zaritskaya, Mark Semenuk, Yun Hee Cho, David W LaFleur, Devanshi Shah, Stephen Ullrich, Giampiero Girolomoni, Cristina Albanesi, and Paul A Moore. Regulatory effect of IFN-κ, a novel type I IFN, on cytokine production by cells of the innate immune system. The Journal of Immunology, 169(9):4822–4830, 2002.

[172] Ramona Hurdayal and Frank Brombacher. Interleukin-4 receptor alpha: from innate to adaptive immunity in murine models of cutaneous leishmaniasis. Frontiers in immunology, 8:1354, 2017.

[173] Yonghong Wang, Bo-Jiang Shen, and Walter Sebald. A mixed-charge pair in human interleukin 4 dominates high-affinity interaction with the receptor α chain. Proceedings of the National Academy of Sciences, 94(5):1657–1662, 1997.

[174] Rosanne Spolski, Peng Li, and Warren J Leonard. Biology and regulation of IL-2: from molecular mechanisms to human therapy. Nature Reviews Immunology, 18(10):648–659, 2018.

[175] Tongtong Zhang, Wei Hu, and Wei Chen. Plasma membrane integrates biophysical and biochemical regulation to trigger immune receptor functions. Frontiers in Immunology, 12:613185, 2021.

[176] Mijo Simunovic and Gregory A Voth. Membrane tension controls the assembly of curvature-generating proteins. Nature communications, 6(1):7219, 2015.

[177] Rarnesh C Tripathi, NS Borisuth, Susmilha P Kolli, and BrendaJ Tripathi. Trabecular cells express receptors that bind TGF-beta 1 and TGF-beta 2: a qualitative and quantitative characterization. Investigative ophthalmology & visual science, 34(1):260–263, 1993.

[178] Adrian Whitty, Natalya Raskin, Dian L Olson, Christopher W Borysenko, Christine M Ambrose, Christopher D Benjamin, and Linda C Burkly. Interaction affinity between cytokine receptor components on the cell surface. Proceedings of the National Academy of Sciences, 95(22):13165–13170, 1998.

[179] Renate M Weber-Nordt, Marco A Meraz, and Robert D Schreiber. Lipopolysaccharide-dependent induction of IL-10 receptor expression on murine fibroblasts. The Journal of Immunology, 153(8):3734–3744, 1994.

[180] Mübeccel Akdis, Alar Aab, Can Altunbulakli, Kursat Azkur, Rita A Costa, Reto Crameri, Su Duan, Thomas Eiwegger, Andrzej Eljaszewicz, Ruth Ferstl, et al. Interleukins (from IL-1 to IL-38), interferons, transforming growth factor β, and tnf-α: Receptors, functions, and roles in diseases. Journal of Allergy and Clinical Immunology, 138(4):984–1010, 2016.

[181] Lucile E Wrenshall and Jeffrey L Platt. Regulation of T cell homeostasis by heparan sulfate-bound IL-2. The Journal of Immunology, 163(7):3793–3800, 1999.

[182] Hugues Lortat-Jacob, Pierre Garrone, Jaques Banchereau, and Jean-Alexis Grimaud. Human interleukin 4 is a glycosaminoglycan-binding protein. Cytokine, 9(2):101–105, 1997.

[183] Shahram Salek-Ardakani, John R Arrand, David Shaw, and Mike Mackett. Heparin and heparan sulfate bind interleukin-10 and modulate its activity. Blood,The Journal of the American Society of Hematology, 96(5):1879–1888, 2000.

[184] Hugues Lortat-Jacob and Jean-Alexis Grimaud. Interferon-γ binds to heparan sulfate by a cluster of amino acids located in the C-terminal part of the molecule. FEBS letters, 280(1):152–154, 1991.

[185] Jonathan Lee, Sheena Wee, Jayantha Gunaratne, RJE Chua, Raymond AA Smith, Ling Ling, David G Fernig, Kunchithapadam Swaminathan, Victor Nurcombe, and Simon M Cool. Structural determinants of heparin-transforming growth factor-β1 interactions and their effects on signaling. Glycobiology, 25(12):1491–1504, 2015.

[186] Rosemary S Mummery and Christopher C Rider. Characterization of the heparin-binding properties of IL-6. The Journal of Immunology, 165(10):5671–5679, 2000.

[187] Maemunah Hasan, Saloua Najjam, Myrtle Y Gordon, Roslyn V Gibbs, and Christopher C Rider. IL-12 is a heparin-binding cytokine. The Journal of Immunology, 162(2):1064–1070, 1999.

[188] Ding Xu and Jeffrey D Esko. Demystifying heparan sulfate-protein interactions. Annual review of biochemistry, 83(1):129–157, 2014.

[189] Stephane Sarrazin, William C Lamanna, and Jeffrey D Esko. Heparan sulfate proteoglycans. Cold Spring Harbor perspectives in biology, 3(7):a004952, 2011.

[190] Michel Léonetti, Adeline Gadzinski, and Gervaise Moine. Cell surface heparan sulfate proteoglycans influence MHC class II-restricted antigen presentation. The Journal of Immunology, 185(7):3847–3856, 2010.

[191] Omai B Garner, Yu Yamaguchi, Jeffrey D Esko, and Vibeke Videm. Small changes in lymphocyte development and activation in mice through tissue-specific alteration of heparan sulphate. Immunology, 125(3):420–429, 2008.

[192] Trini Teixé, Patricia Nieto-Blanco, Ramon Vilella, Pablo Engel, Manuel Reina, and Enric Espel. Syndecan-2 and-4 expressed on activated primary human CD4+ lymphocytes can regulate T cell activation. Molecular immunology, 45(10):2905–2919, 2008.

[193] Michael Brusilovsky, Olga Radinsky, Limor Cohen, Rami Yossef, Avishai Shemesh, Alex Braiman, Ofer Mandelboim, Kerry S Campbell, and Angel Porgador. Regulation of natural cytotoxicity receptors by heparan sulfate proteoglycans in-cis: A lesson from NKp44. European journal of immunology, 45(4):1180–1191, 2015.

[194] Ritu Chakravarti, Vasileia Sapountzi, and Josephine C Adams. Functional role of syndecan-1 cytoplasmic V region in lamellipodial spreading, actin bundling, and cell migration. Molecular biology of the cell, 16(8):3678–3691, 2005.

[195] Mark D Bass, Kirsty A Roach, Mark R Morgan, Zohreh Mostafavi-Pour, Tobias Schoen, Takashi Muramatsu, Ulrike Mayer, Christoph Ballestrem, Joachim P Spatz, and Martin J Humphries. Syndecan-4-dependent Rac1 regulation determines directional migration in response to the extracellular matrix. The Journal of cell biology, 177(3):527–538, 2007.

[196] Yoshio Yamashita, Kenji Oritani, Erina K Miyoshi, Randolph Wall, Merton Bernfield, and Paul W Kincade. Syndecan-4 is expressed by b lineage lymphocytes and can transmit a signal for formation of dendritic processes. The Journal of Immunology, 162(10):5940–5948, 1999.

[197] Yunmin Jung, Inbal Riven, Sara W Feigelson, Elena Kartvelishvily, Kazuo Tohya, Masayuki Miyasaka, Ronen Alon, and Gilad Haran. Three-dimensional localization of T-cell receptors in relation to microvilli using a combination of superresolution microscopies. Proceedings of the National Academy of Sciences, 113(40):E5916–E5924, 2016.

[198] Gediminas Greicius, Lisa Westerberg, Edward J Davey, Eva Buentke, Annika Scheynius, Johan Thyberg, and Eva Severinson. Microvilli structures on B lymphocytes:inducible functional domains? International immunology, 16(2):353–364, 2004.

[199] Tom Frey, Howard R Petty, and Harden M McConnell. Electron microscopic study of natural killer cell-tumor cell conjugates. Proceedings of the National Academy of Sciences, 79(17):5317–5321, 1982.

[200] Sonja Majstoravich, Jinyi Zhang, Susan Nicholson-Dykstra, Stefan Linder, Wilhelm Friedrich, Katherine A Siminovitch, and Henry N Higgs. Lymphocyte microvilli are dynamic, actin-dependent structures that do not require Wiskott-Aldrich syndrome protein (WASp) for their morphology. Blood, 104(5):1396–1403, 2004.

[201] Ron Orbach and Xiaolei Su. Surfing on membrane waves: Microvilli, curved membranes, and immune signaling. Frontiers in Immunology, 11:567890, 2020.

[202] Morgan Huse. Mechanical forces in the immune system. Nature Reviews Immunology, 17(11):679–690, 2017.

[203] Elisaveta Miladinova, Elena Lilkova, Elena Krachmarova, Kristina Malinova, Peicho Petkov, Nevena Ilieva, Genoveva Nacheva, and Leandar Litov. Heparan sulfate facilitates binding of hIFN-γ to its cell-surface receptor hIFNGR1. International Journal of Molecular Sciences, 23(16):9415, 2022.

[204] Wilton Snead, Carl Hayden, Avinash Gadok, Padmini Rangamani, and Jeanne Stachowiak. Membrane fission by protein crowding. Biophysical Journal, 112(3):327a, 2017.

[205] Hao Xing, Huan Liu, Zhengqi Chang, and Ji Zhang. Research progress on the immunological functions of Piezo1 a receptor molecule that responds to mechanical force. International Immunopharmacology, 139:112684, 2024.

[206] Radhakrishnan Gnanasambandam, Chilman Bae, Philip A Gottlieb, and Frederick Sachs. Ionic selectivity and permeation properties of human PIEZO1 channels. PloS one, 10(5):e0125503, 2015.

[207] Pauline Wales, Christian E Schuberth, Roland Aufschnaiter, Johannes Fels, Ireth Garćia-Aguilar, Annette Janning, Christopher P Dlugos, Marco Schäfer-Herte, Christoph Klingner, Mike Wälte, et al. Calcium-mediated actin reset (CaAR) mediates acute cell adaptations. elife, 5:e19850, 2016.

[208] Madhuri S Salker, Nicolas Schierbaum, Nour Alowayed, Yogesh Singh, Andreas F Mack, Christos Stournaras, Tilman E Schäffer, and Florian Lang. LeftyA decreases actin polymerization and stiffness in human endometrial cancer cells. Scientific reports, 6(1):29370, 2016.

[209] Lucas Lamparter and Milos Galic. Cellular membranes, a versatile adaptive composite material. Frontiers in cell and developmental biology, 8:684, 2020.

[210] Cornelis Storm, Jennifer J Pastore, Fred C MacKintosh, Tom C Lubensky, and Paul A Janmey. Nonlinear elasticity in biological gels. Nature, 435(7039):191–194, 2005.

[211] Yan Guan, Xiaonan Shan, Fenni Zhang, Shaopeng Wang, Hong-Yuan Chen, and Nongjian Tao. Kinetics of small molecule interactions with membrane proteins in single cells measured with mechanical amplification. Science Advances, 1(9):e1500633, 2015.

[212] Matthias Pöhnl, Marius FW Trollmann, and Rainer A Böckmann. Nonuniversal impact of cholesterol on membranes mobility, curvature sensing and elasticity. Nature Communications, 14(1):8038, 2023.

[213] Andrew R Houk, Alexandra Jilkine, Cecile O Mejean, Rostislav Boltyanskiy, Eric R Dufresne, Sigurd B Angenent, Steven J Altschuler, Lani F Wu, and Orion D Weiner. Membrane tension maintains cell polarity by confining signals to the leading edge during neutrophil migration. Cell, 148(1):175–188, 2012.

[214] Konrad Pazdrak, Louis Justement, and Rafeul Alam. Mechanism of inhibition of eosinophil activation by transforming growth factor-beta. Inhibition of Lyn, MAP, Jak2 kinases and STAT1 nuclear factor. The Journal of Immunology, 155(9):4454–4458, 1995.

[215] Giulio Preta. New insights into targeting membrane lipids for cancer therapy. Frontiers in cell and developmental biology, 8:571237, 2020.

[216] Claire Gorby, Junel Sotolongo Bellón, Stephan Wilmes, Walid Warda, Elizabeth Pohler, Paul K Fyfe, Adeline Coäni, Christophe Ferrand, Mark R Walter, Suman Mitra, et al. Engineered IL-10 variants elicit potent immunomodulatory effects at low ligand doses. Science signaling, 13(649):eabc0653, 2020.

[217] Ankit Saxena, Sam Khosraviani, Sanjeev Noel, Divya Mohan, Thomas Donner, and Abdel Rahim A Hamad. Interleukin-10 paradox: A potent immunoregulatory cytokine that has been difficult to harness for immunotherapy. Cytokine, 74(1):27–34, 2015.

