## Supplementary materials for "Lymphocytes and monocytes undergo swift suppression of IL-10R, IL-6R, and IL-2Rβγ signaling under high concentrations of different cytokines"

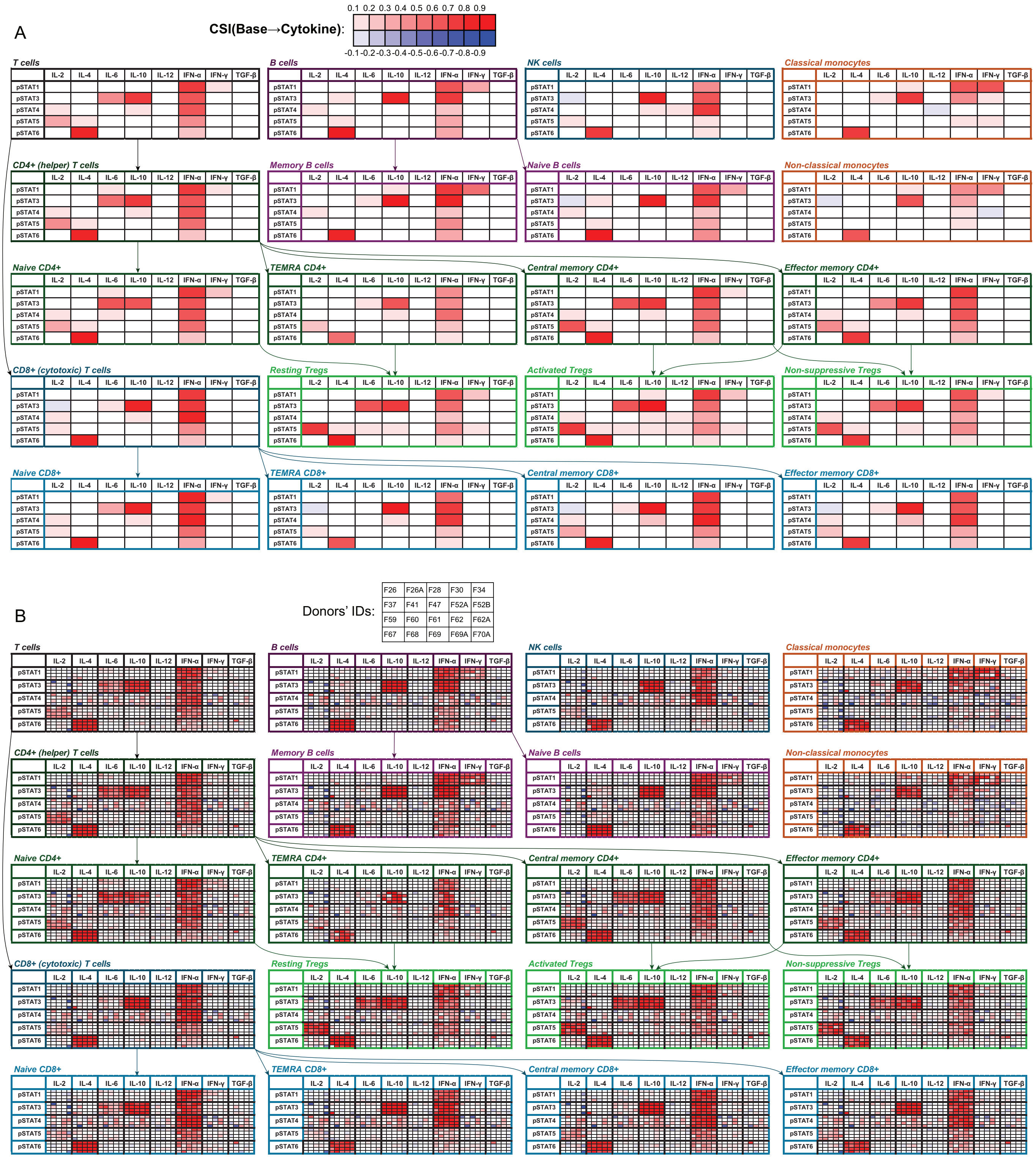


Figure S1: **Outcomes of 15-minute application of single cytokines to immune cells.** A) Aggregate analysis using cells from 19 healthy donors, with equal number of cells sampled from each donor for each setting. B) Individual donor analysis using cells from 20 healthy donors. CSI = Cumulative Shift Index, here and further its definition and the criteria for detecting pSTAT signaling are consistent with those outlined in the main text. Here and further the numbers in the donor IDs correspond to the ages of the respective donors; cell counts for each setting are provided in the supplementary Excel (.xlsx) files. Donor F67's cells are not included in the aggregate analysis.

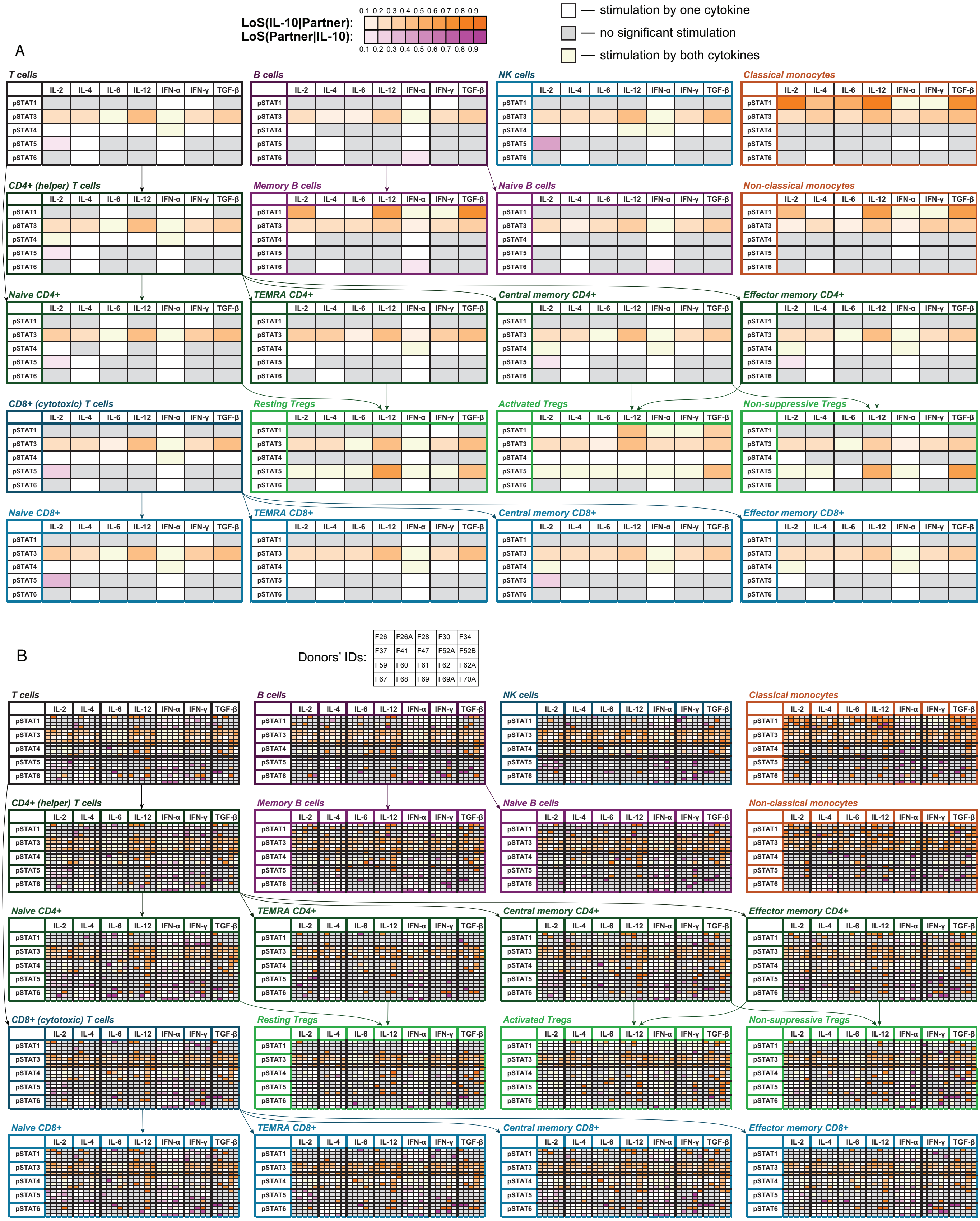


Figure S2: **Outcomes of paired 15-minute application of IL-10 with other cytokines to immune cells.** A) Aggregate analysis using cells from 19 healthy donors, with equal number of cells sampled from each donor for each setting. B) Individual donor analysis using cells from 20 healthy donors. Orange cells denote suppression of IL-10 signaling by other cytokines, purple cells correspond to suppression of other cytokines' signaling by IL-10. Gray cells denote the cases where neither cytokine induces the relevant pSTAT significantly, i.e., both related p-values estimated by the Mann–Whitney U test are $\geq 0.001$. Yellow cells highlight the cases where both cytokines significantly induce the relevant pSTAT, making detection of suppression infeasible. LoS = Level of Suppression, here and further its definition and the criteria for detecting pSTAT signaling are consistent with those outlined in the main text. Donor F67 is not included in the aggregate analysis.

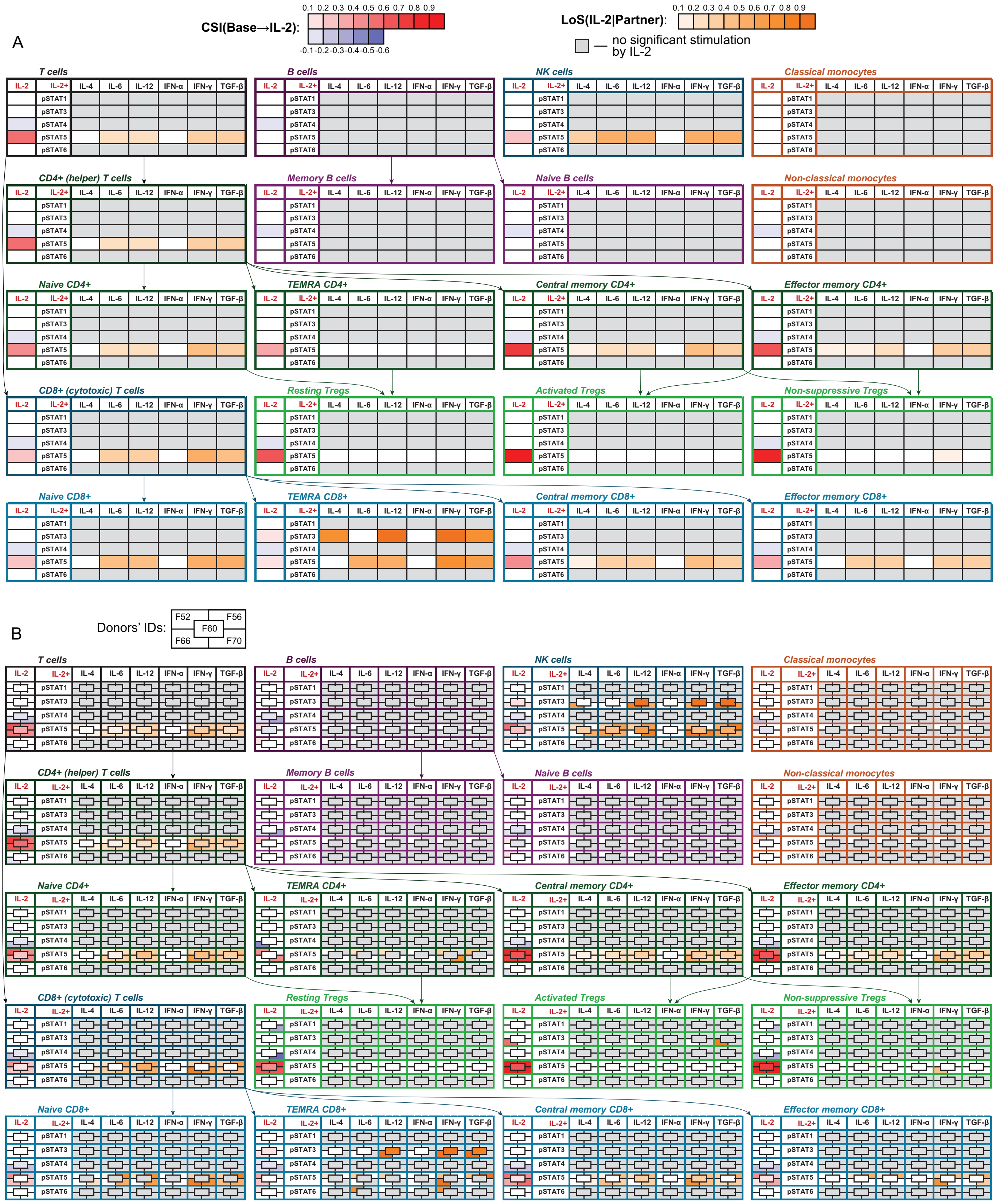

Figure S3: **Outcomes of paired 15-minute application of IL-2 with other cytokines to immune cells.** A) Aggregate analysis using cells from 5 healthy donors, with equal number of cells sampled from each donor for each setting. B) Individual donor analysis using cells from the same 5 donors.

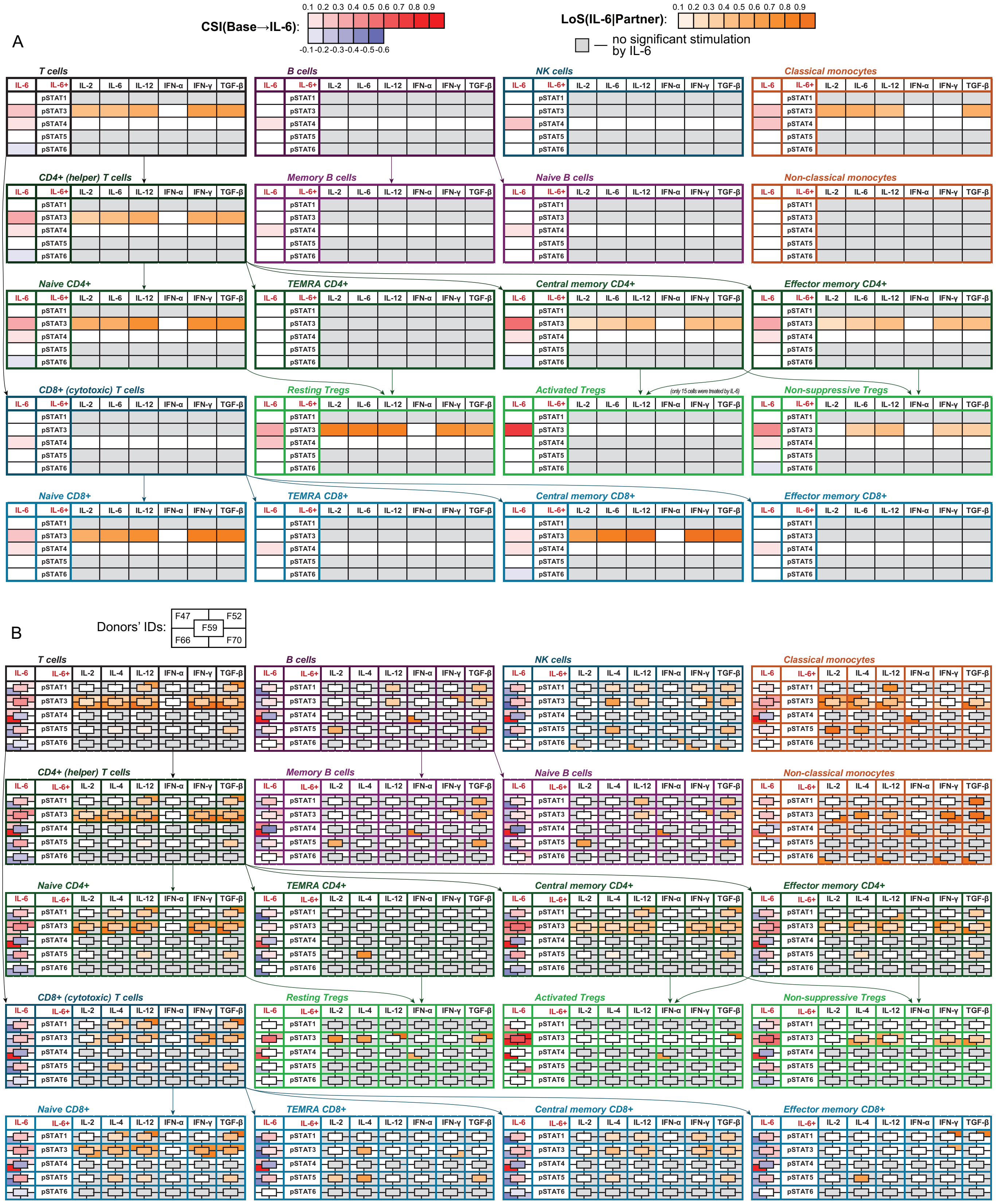


Figure S4: **Outcomes of paired 15-minute application of IL-6 with other cytokines to immune cells.** A) Aggregate analysis using cells from 5 healthy donors, with equal number of cells sampled from each donor for each setting. B) Individual donor analysis using cells from the same 5 donors.

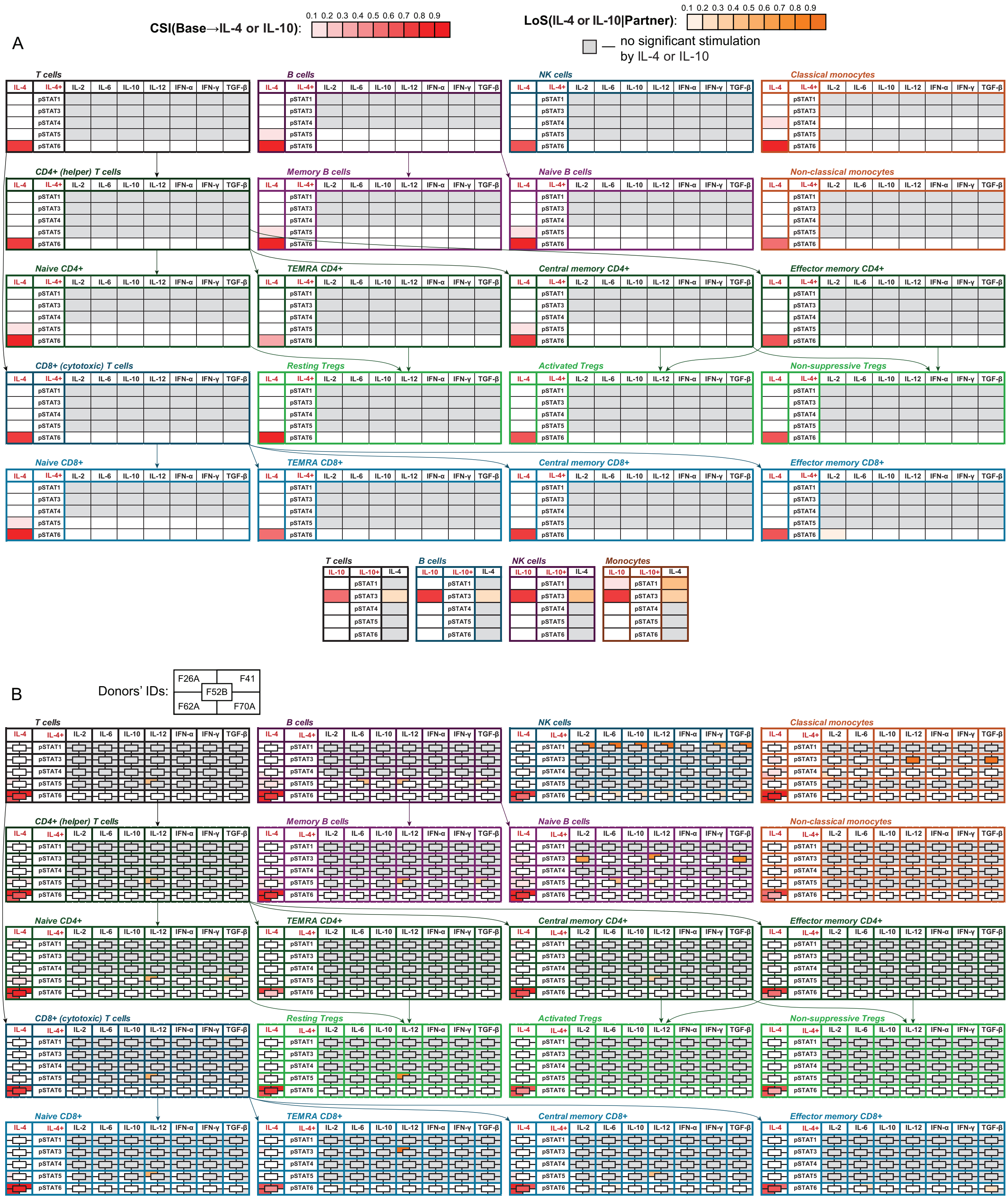


Figure S5: **Outcomes of paired 15-minute application of IL-4 with other cytokines to immune cells.** A) Aggregate analysis using cells from 5 healthy donors, with equal number of cells sampled from each donor for each setting. B) Individual donor analysis using cells from the same 5 donors.

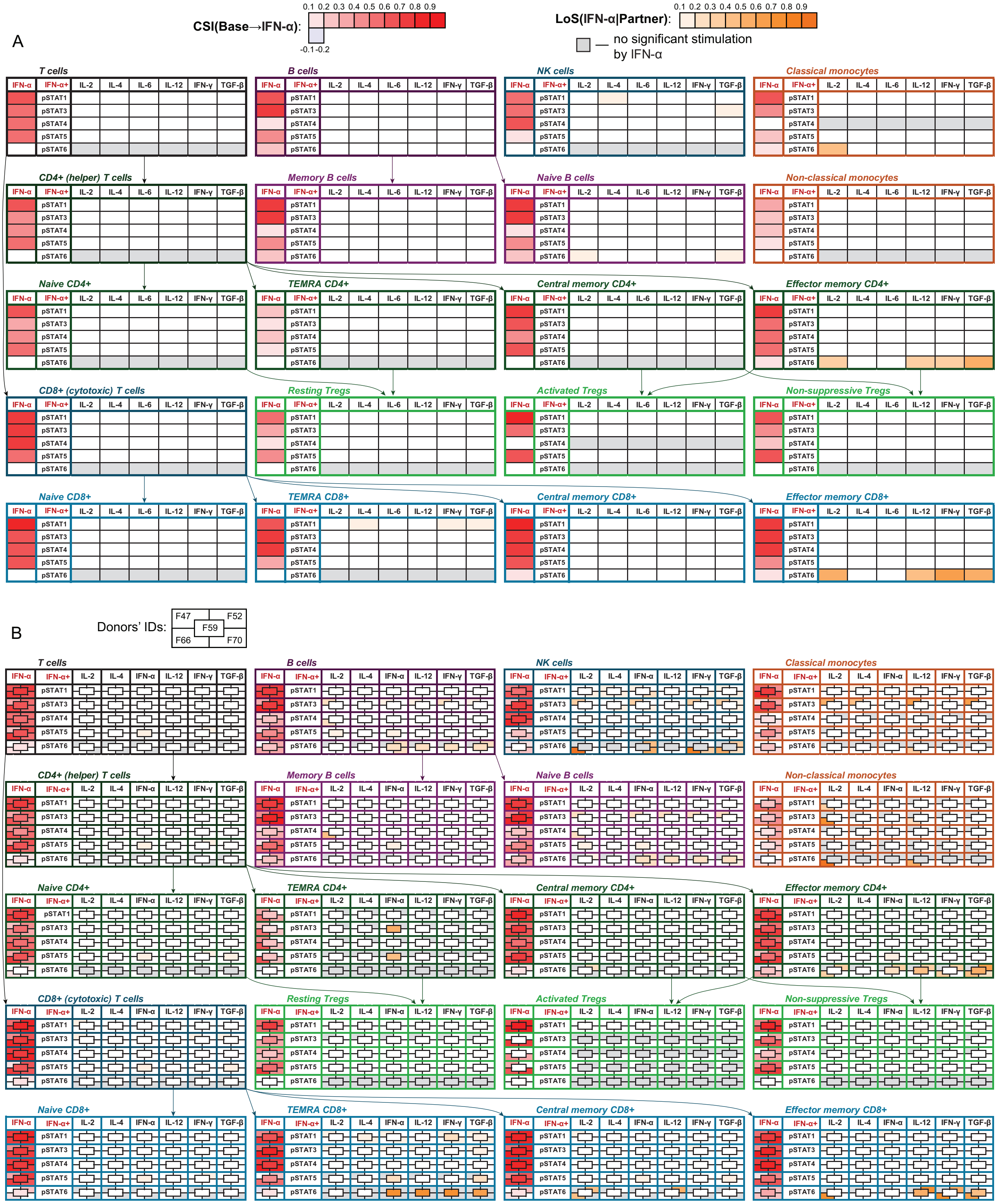


Figure S6: **Outcomes of paired 15-minute application of IFN-α with other cytokines to immune cells.** A) Aggregate analysis using cells from 5 healthy donors, with equal number of cells sampled from each donor for each setting. B) Individual donor analysis using cells from the same 5 donors.

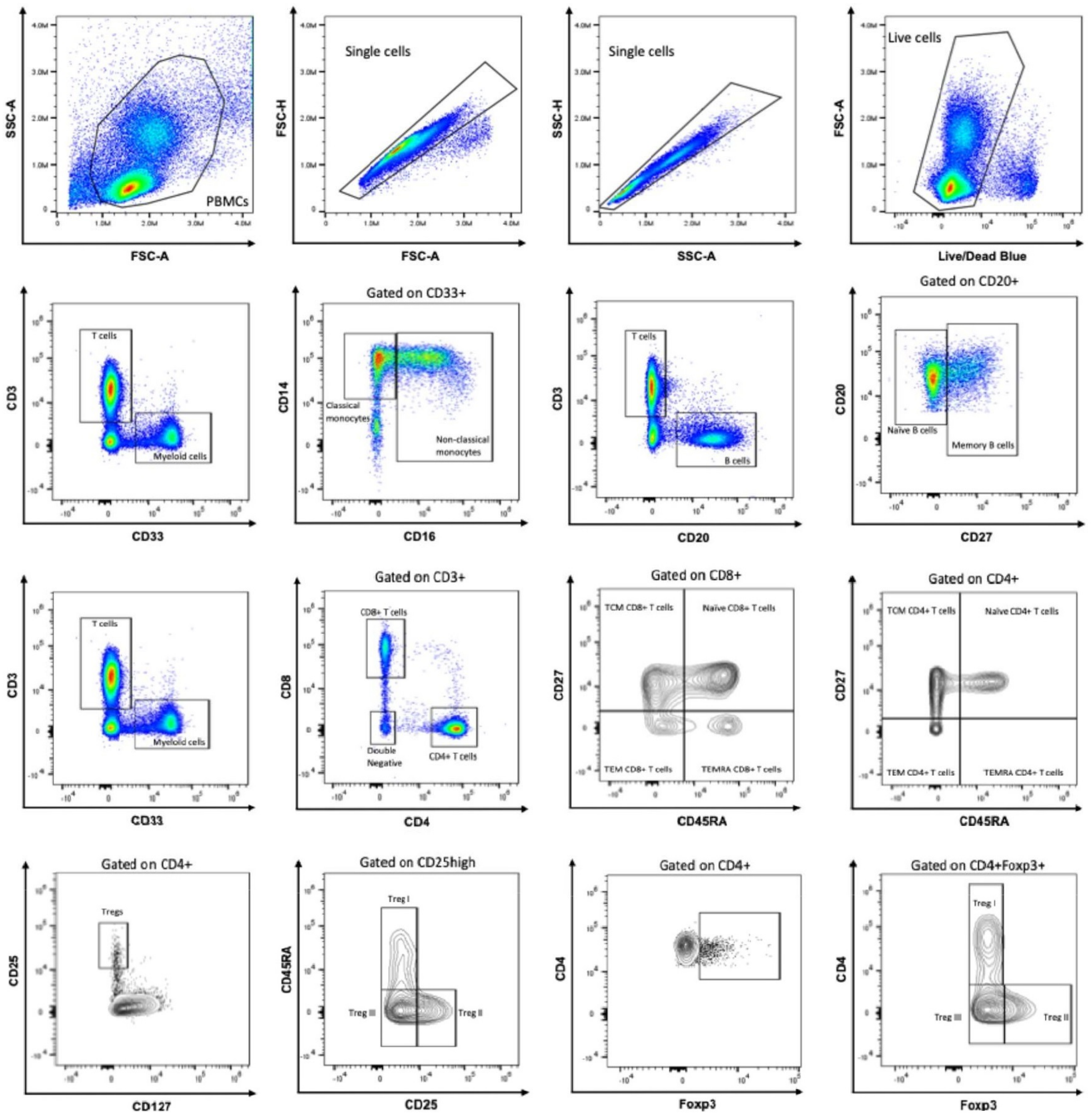


Figure S7: **Gating strategies for identifying cell populations used in this study.** The panels show strategies for identification of PBMCs, single cells, live cells, myeloid cells, monocytes, B cell and T cell subsets.

| **IL-10/STAT3 &** | **IL-4/STAT6** | | **IL-6/STAT3** | | **IFN-$\alpha$/STAT1** | | **IFN-$\alpha$/STAT3** | | **IFN-$\alpha$/STAT4** | | **IFN-$\alpha$/STAT5** | | **IFN-$\alpha$/STAT6** | | **IFN-$\gamma$/STAT1** | | **IL-2/STAT5** | |
|---|---|---|---|---|---|---|---|---|---|---|---|---|---|---|---|---|---|---|
| Cell type | Corr | $p$ | Corr | $p$ | Corr | $p$ | Corr | $p$ | Corr | $p$ | Corr | $p$ | Corr | $p$ | Corr | $p$ | Corr | $p$ |
| Classical monocytes | 0.77 | <0.001 | 0.65 | 0.008 | 0.75 | <0.001 | 0.52 | 0.02 | 0.02 | 0.95 | 0.17 | 0.57 | 0.27 | 0.36 | 0.9 | <0.001 | | |
| Non-classical monocytes | 0.72 | <0.001 | | | 0.92 | <0.001 | 0.59 | 0.007 | | | | | | | 0.79 | <0.001 | | |
| CD16+ NK cells | 0.85 | <0.001 | | | 0.72 | <0.001 | 0.84 | <0.001 | 0.32 | 0.18 | 0.61 | 0.01 | 0.42 | 0.11 | <0.001 | <0.001 | -0.2 | 0.45 |
| Naive CD4+ | 0.33 | 0.16 | 0.86 | <0.001 | 0.59 | 0.007 | 0.81 | <0.001 | 0.46 | 0.04 | 0.61 | 0.005 | 0.17 | 0.48 | 0.5 | 0.03 | -0.02 | 0.92 |
| TEMRA CD4+ | 0.76 | <0.001 | | | 0.83 | <0.001 | 0.9 | <0.001 | 0.9 | <0.001 | 0.02 | 0.95 | <0.001 | <0.001 | | | -0.45 | 0.13 |
| TCM CD4+ | 0.57 | 0.008 | 0.57 | 0.008 | 0.38 | 0.1 | 0.52 | 0.02 | 0.24 | 0.31 | 0.32 | 0.17 | 0.17 | 0.48 | 0.29 | 0.25 | -0.07 | 0.76 |
| TEM CD4+ | 0.33 | 0.16 | -0.29 | 0.21 | 0.31 | 0.19 | 0.39 | 0.09 | 0.47 | 0.03 | -0.05 | 0.84 | 0.02 | 0.95 | | | -0.53 | 0.02 |
| Naive CD8+ | -0.02 | 0.94 | 0.23 | 0.33 | 0.01 | 0.96 | 0.64 | 0.002 | 0.63 | 0.003 | 0.07 | 0.78 | -0.01 | 0.96 | 0.12 | 0.66 | 0.21 | 0.41 |
| TEMRA CD8+ | 0.18 | 0.46 | | | 0.21 | 0.39 | 0.34 | 0.15 | 0.27 | 0.25 | 0.14 | 0.59 | -0.17 | 0.55 | | | -0.04 | 0.89 |
| TCM CD8+ | -0.11 | 0.66 | 0.1 | 0.68 | -0.11 | 0.63 | 0.01 | 0.95 | 0.4 | 0.08 | 0.02 | 0.95 | -0.03 | 0.89 | | | -0.1 | 0.72 |
| TEM CD8+ | 0.32 | 0.16 | | | 0.31 | 0.19 | 0.58 | 0.008 | 0.54 | 0.01 | 0.29 | 0.22 | 0.29 | 0.26 | <0.001 | <0.001 | 0.06 | 0.83 |
| Naive B cells | 0.3 | 0.2 | | | 0.35 | 0.13 | 0.45 | 0.05 | -0.1 | 0.69 | 0.11 | 0.63 | -0.1 | 0.67 | -0.07 | 0.76 | | |
| Memory B cells | 0.27 | 0.24 | | | 0.15 | 0.53 | 0.1 | 0.67 | -0.37 | 0.15 | -0.05 | 0.85 | -0.06 | 0.83 | 0.45 | 0.05 | | |

Table S1: **Spearman's rank correlation coefficients for individual cytokine signaling.** Coefficients are indicated for correlation between IL-10/STAT3 and other signaling axes in different types of lymphocytes and monocytes.

| | IL-10 | IL-6 | IFN-$\alpha$ | IFN-$\gamma$ | IL-4 | IL-2 |
|---|---|---|---|---|---|---|
| Drives immune system to | **Suppressed activity** | **Broad response skewed to type 3** *extracellular bacteria, fungi* | **Broad response skewed to type 1** | **Type 1 response** *intracellular microbes, tumors* | **Type 2 response** *parasites, allergens* | **Enhancement of lymphocyte activity** |
| Main producers | Tregs, Th1, Th2, Th17 cells, DCs[1] | Macrophages, B and T cells, DCs[2] | Macrophages, DCs[3] | NK, NKT, Th1, CD8+ effector cells[4] | Th2 cells, mast cells, basophils, eosinophils[5] | CD4+ T cells at the onset of activation[6] |
| *Cell type* | *Directly induced effects at normal physiological concentrations* | | | | | |
| CD8+ T cells | Reduced antigen sensitivity[7] | Differentiation into Foxp3+[8] and IL-21-producing B helper CD8+ T cells[9] | Enhanced expansion, IFN-$\gamma$ secretion, cytotoxicity[10]; Co-stimulation of IL-4 signaling[11] | **Enhanced cytotoxicity and motility**[12] | Long-term survival of memory cells[13] | Proliferation; Differentiation of primed cells into effector cells[14] |
| CD4+ T cells | Suppressed proliferation and IL-2, IFN-$\gamma$, IL-4 secretion[15] | IL-4 secretion[16]; Th17 differentiation (with TGF-$\beta$)[17] | Enhanced IFN-$\gamma$ secretion[18]; Co-stimulation of IL-4 signaling[11] | Upregulation of IL-12R$\beta$2 for Th1 differentiation[19]; Enhanced survival[20] | Th2 differentiation[21] | Differentiation[14]; Maintenance of a functional Treg population[22] |
| B cells | Apoptosis upon activation[23] | Proliferation; IgM production[24] | Lowered activation threshold[25] | Proliferation; IgG2 production[24] | **Proliferation; IgG1 and IgE production**[24] | Proliferation, differentiation[26] |
| NK cells | Suppressed IFN-$\gamma$ secretion[27] | Suppressed cytotoxicity[28] | Activation[29] | **[enhanced MHC-I expression on healthy cells inhibiting lysis by NK cells]**[30] | Suppressed IL-2-driven activation and proliferation[31] | Proliferation and partial activation[32]; Enhanced cytotoxicity[33] |
| Mono-cytes | Suppressed APC function and cytokine secretion[34] | Chemokine secretion[35] | Enhanced APC function[36] | Differentiation into phagocytic macrophages[37] | Suppressed secretion of IL-1 and TNF-$\alpha$[38] | [no effect at physiological levels][39] |
| Macro-phages | Suppressed cytokine secretion[40] | Regulatory polarization and proliferation[41] | Enhanced ROS release[42] | **Enhanced activation and cytotoxicity**[43] | Regulatory polarization[44] | Slight reduction of activation threshold[45] |
| DCs | Suppressed maturation[46] and chemokine secretion[47] | Suppressed maturation[48] | Activation[49] | Maturation for efficient APC function[50] | Suppression of response to IFN-$\alpha$[51] | Secrete IL-2 upon pattern recognition receptors stimulation[52] |
| Neutro-phils | Suppressed TNF-$\alpha$ and chemokine secretion[53] | **Secretion of TNF-$\alpha$ and IL-8 mediated by release of serum amyloid A**[54] | Enhanced TNF-$\alpha$ and ROS release[55] | Promotion of extracellular trap formation for bacteria elimination[56] | Suppressed expansion and migration[57] | [no effect at physiological levels][58] |
| Eosino-phils | Suppressed activation and ROS secretion[59] | [can produce and store IL-6][60] | Suppressed cytotoxicity against parasites[61] | Enhanced ROS release, chemotaxis, and survival[62] | **Enhanced migration**[63] | Chemotaxis towards IL-2[64] |

Table S2: **Direct effects of cytokines.** This framework supposedly drives maintenance of IFN-$\alpha$, IFN-$\gamma$, and IL-4 signaling, and suppression of IL-10, IL-6, and (transiently) IL-2 signaling for enhanced efficiency of immune response. Bold text indicates crucial functions directly associated with killing of pathogens. AICD = activation-induced cell death, APC = antigen-presenting cell, DC = dendritic cell, ROS = reactive oxygen species.

Table S3: **Pro-inflammatory effects attributed to IL-10.** These effects likely resulted from non-specific immune cell reaction to high cytokine concentrations. i.p. = intraperitoneally, s.c. = subcutaneously, i.v. = intravenously, PBMC = peripheral blood mononuclear cells, LPS = lipopolysaccharide.

*In vitro studies*

| Cell type | Effects attributed to IL-10 | Treatment | Comments | Ref. |
|---|---|---|---|---|
| CD8+ T cells | Enhanced secretion of granzyme and perforin in IFN-$\gamma^{-/-}$ cells | 1-1,000 ng/mL per $10^6$ cells for 24 h before activation | Slight enhancement of granzyme and perforin secretion under 1 ng/mL of IL-10, strong enhancement under 10-1,000 ng/mL of IL-10. | 65 |
| CD8+, CD4+ T cells | Enhanced survival with lower proliferation rate, but higher viability | 1 µg/mL per $5 \cdot 10^4$ cells for 0-14 days before activation | Cells from ulcerative colitis remission patients, in whom higher serum IL-10 levels were associated with shorter remission. IL-10 added 3 days pre-stimulation was the most effective at increasing the live cell numbers. | 66 |
| CD4+ T cells | Enhanced production of IL-2, IFN-$\gamma$, IL-4, TNF-$\alpha$ after activation | 20 ng/mL per $10^6$ cells for 24 h before activation | Different cell clones required different incubation period to show similar effects (24-48 h). In contrast, when IL-10 was added only during activation, it inhibited IL-2, but not IFN-$\gamma$, IL-4, TNF-$\alpha$ production. | 67 |
| B cells | Enhanced proliferation of activated cells | 0.5-1,000 ng/mL per $2.5 \cdot 10^4$ cells/100 µL for 9 days | Modest increase of proliferation rate at 0.5 ng/mL of vIL-10, saturation at ~100 ng/mL of vIL-10 and a ~1000 ng/mL of hIL-10. Proliferation slowed down at day 7. Proliferation rate increased significantly under co-application of ~5 ng of IL-4. | 68 |
| B cells | Enhanced IgM, IgG, IgA production | 10 ng/mL per $2 \cdot 10^5$ cells/0.5 mL in presence of activating signals for 10 days | Addition of ~0.1 ng/mL of IL-4 resulted in "a 50% decrease of the IL-10-induced IgM, IgG, and IgA production; maximal inhibition (75%) was obtained with IL-4 at ~1 ng/mL. Addition of IL-10 only weakly affected IL-4-induced IgE production." | 68 |
| B cells | Enhanced differentiation into plasma cells | 100 ng/mL per $2 \cdot 10^6$ cells/0.2 mL for 3 days | Treatment was performed in presence of plasma cell formation-driving signals. "Addition of IL-2 to the IL-10 and CD70-transfect activation system greatly induced differentiation into plasma cells." | 69 |
| NK cells | Increased IFN-$\gamma$ production and cytotoxicity | 50 ng/mL per $\lesssim 10^6$ cells for 16 hours | Pre-incubation with IL-10 was followed by activation by target cells. The effects were enhanced upon the increase of IL-10 concentration from 5 to 50 ng/mL and were saturated at 100 ng/mL of IL-10. | 70 |
| NK cells | Enhanced cytotoxicity and mRNA levels of cytotoxicity-related and activation genes | 30 ng/mL per $10^5$ cells for 4 h | No effect on proliferation and migration. "The choice of IL-10 dose was derived from dose–effect experiments, which showed a maximum number of genes changing their transcriptional levels at IL-10 concentrations ≥30 ng/mL." | 71 |
| PBMC | Stimulation of cytolytic activity | 3-75 ng/mL per $10^6$ cells for 3 days before cytotoxicity assays | "Induction of cytolytic activities by IL-10 was neutralized by anti-IL-10 monoclonal antibodies but not by antibodies against IFN-$\gamma$ or TNF-$\alpha$. Co-incubation of PBMCs with IL-10 and IL-2 or IL-10 and IFN-$\alpha$ augmented cytolytic activity, in particular at lower effector-to-target ratios." | 72 |

*In vivo studies*

| Cell type | Effects attributed to IL-10 | Dose or serum level (SL) | Comments | Ref. |
|---|---|---|---|---|
| CD8+ T cells | Activation and expansion of tumor-resident cells | ~0.25-2 µg/mL (initial SL) | C57BL/6 mice with established PDV6 tumors were injected with 10 µg of IL-10 minicircle. | 73 |

| | | | | |
|---|---|---|---|---|
| CD8+ T cells | Infiltration and activation of intratumoral cells, enhanced secretion of IFN-$\gamma$ and granzymes | 0.1-0.5 mg/kg s.c. daily or twice daily for 2-4 weeks | Pegylated IL-10 was used. "Most of the areas in IL-10-treated tumors showed a significantly higher expression of MHC class II." | 65 |
| CD8+ T cells | Enhancement of antigen-specific proliferation | 40 µg/mouse i.p. daily for 4 days | Injection of IL-10 "just after a booster vaccine significantly enhanced antitumor immunity and vaccine efficacy." "Administration of IL-10 before, or soon after, peptide-pulsed primary dendritic cell immunization resulted in immune suppression and enhanced tumor progression." | 74 |
| CD8+, CD4+ T cells | Accelerated lethality of autoimmune graft-versus-host disease | 0.01-30 µg i.p. twice daily for 2 weeks | The group receiving 1 ng doses of IL-10, in contrast, showed increased survival, while 0.1 ng doses did not affect survival. | 75 |
| NK cells | Abundant localization in tumor metastases | 1-150 µg/kg i.p. | Inhibition of metastases was observed in mice deficient in T or B cells, but not in mice deficient in NK cells. | 76 |
| T and NK cells | Enhanced cytotoxic activity | 20-60 µg i.p. for 7 days | "Rejection of tumors, delaying tumor outgrowth or resulting in complete cure." "Cured mice were immune to subsequent rechallenge with 10-fold higher inoculation with the same, but not a different, tumor." | 77 |

*Clinical studies*

| **Setting** | **Effects attributed to IL-10** | **Dose** | **Comments** | **Ref.** |
|---|---|---|---|---|
| Crohn's disease | Upregulation of IFN-$\gamma$ production | 20 µg/kg s.c. daily for 4 weeks | Increase in serum neopterin (produced by monocytes and macrophages activated by IFN-$\gamma$) after 2 weeks. No clinical efficacy in patients with moderate or active disease. Patients with moderate disease receiving 5 µg/kg had remission. Higher doses were less effective. | 78,79 |
| Solid tumors | CD8+ T cell-mediated immune activation, elevation of IFN-$\gamma$ and Granzyme B | 20 µg/kg of PEGylated IL-10 daily s.c. for 4 weeks | Objective tumor responses in 4/15 patients with intermediate- to poor-risk renal cell cancer, without autoimmune toxicities. 15/41 patients with advanced disease had durable disease stabilization. PEGylated IL-10 was elevated above 1 ng/mL in serum. | 80 |
| Healthy subjects | Enhanced activation of CD8+ T and NK cells, enhanced LPS-induced IFN-$\gamma$ release | 25 µg/kg i.v. | LPS (4 ng/kg) was administered just after or 1 h before IL-10. IL-10 inhibited or not influenced the production of IFN-$\gamma$-inducing cytokines. | 81 |
| Healthy subjects | Transient increase of circulating neutrophils and monocytes in a dose-dependent manner | 5-100 µg/kg i.v. | Transient decrease of circulating lymphocytes. Mild to moderate flu-like syndrome at 100 µg/kg. "Inhibition of cytokine synthesis by peripheral blood cells stimulated *ex vivo* with bacterial LPS." | 82 |

### References


[1] Valentina Carlini, Douglas M Noonan, Eslam Abdalalem, Delia Goletti, Clementina Sansone, Luana Calabrone, and Adriana Albini. The multifaceted nature of IL-10: regulation, role in immunological homeostasis and its relevance to cancer, COVID-19 and post-COVID conditions. *Frontiers in immunology*, 14:1161067, 2023.

[2] Melissa Swiecki and Marco Colonna. Type I interferons: diversity of sources, production pathways and effects on immune responses. *Current opinion in virology*, 1(6):463–475, 2011.

[3] Olof Berggren, Andrei Alexsson, David L Morris, Karolina Tandre, Gert Weber, Timothy J Vyse, Ann-Christine Syvänen, Lars Rönnblom, and Maija-Leena Eloranta. IFN-$\alpha$ production by plasmacytoid dendritic cell associations with polymorphisms in gene loci related to autoimmune and inflammatory diseases. *Human Molecular Genetics*, 24(12):3571–3581, 2015.

[4] Jamie R Schoenborn and Christopher B Wilson. Regulation of interferon-$\gamma$ during innate and adaptive immune responses. *Advances in immunology*, 96:41–101, 2007.

[5] P Sachin, S Gadani, J Cronk, G Norris, and J Kipnis. Interleukin-4: a cytokine to remember. *The Journal of Immunology*, 189:4213–4421, 2012.

[6] Wei Liao, Jian-Xin Lin, and Warren J Leonard. IL-2 family cytokines:new insights into the complex roles of IL-2 as a broad regulator of T helper cell differentiation. *Current opinion in immunology*, 23(5):598–604, 2011.

[7] Logan K Smith, Giselle M Boukhaled, Stephanie A Condotta, Sabrina Mazouz, Jenna J Guthmiller, Rahul Vijay, Noah S Butler, Julie Bruneau, Naglaa H Shoukry, Connie M Krawczyk, et al. Interleukin-10 directly inhibits CD8+ T cell function by enhancing N-glycan branching to decrease antigen sensitivity. *Immunity*, 48(2):299–312, 2018.

[8] Takayuki Nakagawa, Mineko Tsuruoka, Hideki Ogura, Yuko Okuyama, Yasunobu Arima, Toshio Hirano, and Masaaki Murakami. IL-6 positively regulates Foxp3+CD8+ T cells in vivo. *International immunology*, 22(2):129–139, 2010.

[9] Rui Yang, April R Masters, Karen A Fortner, Devin P Champagne, Natalia Yanguas-Casás, Daniel J Silberger, Casey T Weaver, Laura Haynes, and Mercedes Rincon. IL-6 promotes the differentiation of a subset of naive CD8+T cells into IL-21-producing B helper CD8+ T cells. *Journal of Experimental Medicine*, 213(11):2281–2291, 2016.

[10] Julie M Curtsinger, Javier O Valenzuela, Pujya Agarwal, Debra Lins, and Matthew F Mescher. Cutting edge: type I IFNs provide a third signal to CD8 T cells to stimulate clonal expansion and differentiation. *The Journal of Immunology*, 174(8):4465–4469, 2005.

[11] Karsten Wessel Eriksen, Viveca Horst Sommer, Anders Woetmann, Anette Bødker Rasmussen, Christine Brender, Arne Svejgaard, Søren Skov, Carsten Geisler, and Niels Ødum. Bi-phasic effect of interferon (IFN)-$\alpha$: IFN-$\alpha$ up-and down-regulates interleukin-4 signaling in human T cells. *Journal of Biological Chemistry*, 279(1):169–176, 2004.

[12] Purnima Bhat, Graham Leggatt, Nigel Waterhouse, and Ian H Frazer. Interferon-$\gamma$ derived from cytotoxic lymphocytes directly enhances their motility and cytotoxicity. *Cell death & disease*, 8(6):e2836–e2836, 2017.

[13] Li-Rung Huang, Fen-Ling Chen, Yi-Ting Chen, Ya-Min Lin, and John T Kung. Potent induction of long-term CD8+ T cell memory by short-term IL-4 exposure during t cell receptor stimulation. *Proceedings of the National Academy of Sciences*, 97(7):3406–3411, 2000.

[14] Onur Boyman and Jonathan Sprent. The role of interleukin-2 during homeostasis and activation of the immune system. *Nature Reviews Immunology*, 12(3):180–190, 2012.

[15] Kevin N Couper, Daniel G Blount, and Eleanor M Riley. IL-10:the master regulator of immunity to infection. *The Journal of Immunology*, 180(9):5771–5777, 2008.

[16] Sean Diehl, Chi-Wing Chow, Linda Weiss, Alois Palmetshofer, Thomas Twardzik, Laura Rounds, Edgar Serfling, Roger J Davis, Juan Anguita, and Mercedes Rincón. Induction of NFATc2 expression by interleukin 6 promotes T helper type 2 differentiation. *The Journal of experimental medicine*, 196(1):39–49, 2002.

[17] Song Guo Zheng, Juhua Wang, and David A Horwitz. Cutting edge: Foxp3+CD4+CD25+ regulatory T cells induced by IL-2 and TGF-$\beta$ are resistant to Th17 conversion by IL-6. *The Journal of Immunology*, 180(11):7112–7116, 2008.

[18] Volker Brinkmann, Thomas Geiger, Sefik Alkan, and Christoph H Heusser. Interferon alpha increases the frequency of interferon gamma-producing human CD4+ T cells. *The Journal of experimental medicine*, 178(5):1655–1663, 1993.

[19] Ronald B Smeltz, June Chen, Rolf Ehrhardt, and Ethan M Shevach. Role of IFN-$\gamma$ in Th1 differentiation: IFN-$\gamma$ regulates IL-18R$\alpha$ expression by preventing the negative effects of IL-4 and by inducing/maintaining IL-12 receptor $\beta$2 expression. *The Journal of Immunology*, 168(12):6165–6172, 2002.

[54] Cristiane Jaciara Furlaneto and Ana Campa. A novel function of serum amyloid A: a potent stimulus for the release of tumor necrosis factor-$\alpha$, interleukin-1$\beta$,and interleukin-8 by human blood neutrophil. *Biochemical and biophysical research communications*, 268(2):405–408, 2000.
